# Nutrient Availability Modulates Beneficial Effect of Bacterial Community Volatiles and Contact-Dependent Interactions Differently

**DOI:** 10.64898/2026.07.12.738021

**Authors:** Gözde Merve Türksoy, Jannis Stollenwerk, Miroslav Berka, Martin Černý, Stanislav Kopriva

**Affiliations:** Max Planck Institute for Plant Breeding Research, Carl-von-Linne-Weg 10, 50829, Cologne, Germany; Institute for Plant Sciences, Cluster of Excellence on Plant Sciences (CEPLAS), University of Cologne, 50674 Cologne, Germany; Germany; Department of Molecular Biology and Radiobiology, Faculty of AgriSciences, Mendel University in Brno, Brno, Czech Republic

**Keywords:** Plant-Microbe Interactions, bacterial community VOCs, contact and contact independent interactions, nutrient homeostasis, nitrate, sulfate, phosphate, nutrient deficiency, regulatory networks, root exudates

## Abstract

Plant growth–promoting bacteria enhance plant performance, yet how different modes of plant–microbe interaction shape nutrient-specific host responses remains poorly understood. In particular, it is unclear how direct bacterial contact and volatile-mediated interactions originating from the same bacterial community differentially regulate plant nutrient acquisition pathways. Here, we investigated how a 16-member synthetic bacterial community (16SC) affects plant growth, nutrient status, signaling, and metabolite profiles under full nutrient supply as well as nitrogen (N), sulfur (S), and phosphorus (P) limitation in *Arabidopsis thaliana.* We show that volatile organic compounds (VOCs) emitted by the 16SC promote shoot growth under nitrate limitation and full nutrient conditions, whereas this growth promotion is lost under sulfur- and phosphorus-limiting conditions. In contrast, direct interaction (DBC) between plants and the 16SC abolishes growth promotion under all three nutrient-limiting conditions. These nutrient-dependent phenotypes correlate with distinct regulation of nutrient transporters and key transcriptional regulators involved in N (NRT1;1 / NLP7), S (SULTR1;2 / SLIM1/EIL3), and P (PHO2 / PHR1) signaling pathways. Genetic analyses using nutrient transporter mutants revealed that VOC-induced growth promotion requires functional NRT1;1 and SULTR1;2 transporters, whereas growth promotion mediated by direct bacterial contact is retained in the corresponding mutants. This uncoupling of VOC- and contact-dependent effects indicates that distinct host regulatory pathways underlie bacterial community growth promotion depending on the interaction mode. Together, our findings demonstrate that bacterial community–mediated plant growth promotion is strongly shaped by nutrient context and interaction mode, and that volatile-mediated and contact-dependent mechanisms engage separable host nutrient regulatory networks.

## Introduction

Plants require essential macronutrients such as nitrogen (N), phosphorus (P), and sulfur (S) for growth. However, large portions of these elements exist in forms that plants cannot directly absorb. Microorganisms play an indispensable role in supporting plant nutrition and growth (Griffin et al., 2024), facilitating nutrient acquisition and improving plant health (Trivedi et al., 2020). Plant–microbe symbioses significantly enhance nutrient acquisition. Key mechanisms include rhizobacterial nitrogen fixation in legumes (Rahimlou et al., 2021) and mycorrhiza-mediated mobilization of phosphate and sulfate (Gutjahr & Parniske, 2013; Shi et al., 2026; Wang et al., 2017). Plant growth–promoting rhizobacteria (PGPR) represent another important group of beneficial microorganisms. These bacteria colonize the rhizosphere and influence plant growth through multiple mechanisms, including the solubilization of mineral nutrients, the production of siderophores that chelate iron, and the secretion of phytohormones that stimulate root growth (Bulgarelli et al., 2013; Lugtenberg & Kamilova, 2009). Numerous studies have demonstrated that PGPR can increase the bioavailability of nitrogen, phosphorus, sulfur and other micronutrients, ultimately improving plant growth and yield (Lugtenberg & Kamilova, 2009). However, many studies focus on the growth-promoting effects of single bacterial strains, whereas in natural environments bacteria exist within complex microbial communities. Therefore, studies that address this complexity using synthetic communities (SynComs) or soil-based systems are needed.

In addition to direct colonization of plant roots, microorganisms can influence plant physiology through the emission of volatile organic compounds (VOCs). VOCs are low-molecular-weight metabolites characterized by high vapor pressure and low boiling points, enabling them to diffuse through air and soil matrices (Schulz & Dickschat, 2007; Weisskopf et al., 2021). Some VOCs, such as dimethyl disulfide (DMDS) or indole, are produced by multiple microbial taxa, whereas others are specific to particular bacterial groups or species. Increasing evidence indicates that bacterial VOCs can promote plant growth by modulating phytohormone signaling (Bailly et al., 2014; Tahir et al., 2017; H. Zhang et al., 2007, 2009), enhancing nutrient acquisition (del Carmen Orozco-Mosqueda et al., 2013; Meldau et al., 2013; H. Zhang et al., 2009) or acting as antimicrobial agents that protect plants against pathogens and herbivores (Aziz et al., 2016; Garbeva & Weisskopf, 2020; Pieterse et al., 2014). Well-characterized examples include acetoin emitted by *Bacillus* species (Ryu et al., 2003) and dimethyl disulfide produced by *Bacillus* spp. (Meldau et al., 2013). However, whether microbial VOCs influence plant nutrient acquisition and metabolism remains poorly understood.

Here, we used a defined 16-member bacterial synthetic community (16SC) derived from the *Arabidopsis thaliana* root microbiome, representing 16 different bacterial families (Wippel et al., 2021), to investigate how different microbial communication modes influence plant nutrient responses. Both direct bacterial contact (DBC) and VOCs-mediated interactions from this community have previously been shown to promote plant growth (Türksoy et al., 2025), yet the mechanisms underlying these effects remain unclear. We compared how macronutrients availability modulate the effects of direct microbial colonization and VOCs-mediated signaling from the same SynCom on *A. thaliana* growth and nutrient status. Plant growth was quantified by measuring fresh weight, and nutrient status was assessed by determining the accumulation of major inorganic anions, including nitrate (NO₃⁻), phosphate (PO₄³⁻), and sulfate (SO₄²⁻). To dissect the involvement of nutrient signaling and transport pathways, we analyzed mutants impaired in nitrogen, phosphorus, and sulfur responses, including *nrt1;1* and *nlp7* (nitrogen transport and signaling), *phr1* and *pho2* (phosphorus signaling), and *sultr1;2* and *eil3* (sulfur transport and regulation), under both VOC and DBC treatments. Finally, we characterized metabolite profiles in roots, shoots, and root exudates to evaluate how microbial interaction modes shape plant metabolic responses under both nutrient-sufficient and nutrient-deficient conditions.

## Results

### Bacterial community VOC-Mediated Growth Promotion Is Affected by Nutrient Availability

While many studies on bacterial volatile-mediated growth promotion have documented transcriptomic and metabolomic shifts in the plants, a significant knowledge gap remains regarding how these volatiles modulate plant nutrient status. Particularly, the influence of varying nutrient availability on the volatile-plant interaction is poorly understood. To evaluate the influence of nutrients on 16SC VOC-mediated plant growth promotion (PGP), we tested the community’s effect under limiting conditions of low N, low P, and low S. We included low S, because in the VOC bouquet of the 16SC community, numerous sulfur-containing volatiles were detected, most notably DMDS (Türksoy et al., 2025). While the 16SC-mediated growth promotion was abolished under low S and low P, it was maintained under low N (Figure 1A). We further characterized the plant’s internal nutrient status using ion chromatography. The nutrient limitations did not substantially affect growth of the plants and led to decreases in the foliar levels of the corresponding nutrients under mock conditions (Figure 1). Under PGP conditions (full media and low N), we detected a significant increase in shoot nitrate levels, which was accompanied by a concomitant decrease in phosphate and sulfate concentrations (Figure 1B, C, D). Conversely, under low P and low S, where growth promotion was lost, no significant changes in internal nitrate or phosphate levels were observed. Collectively, these results demonstrate that higher shoot nitrate accumulation strongly correlates with VOC-induced biomass increase, suggesting that nitrate signaling or uptake might be connected to the 16SC volatile response.

**Figure 1.**
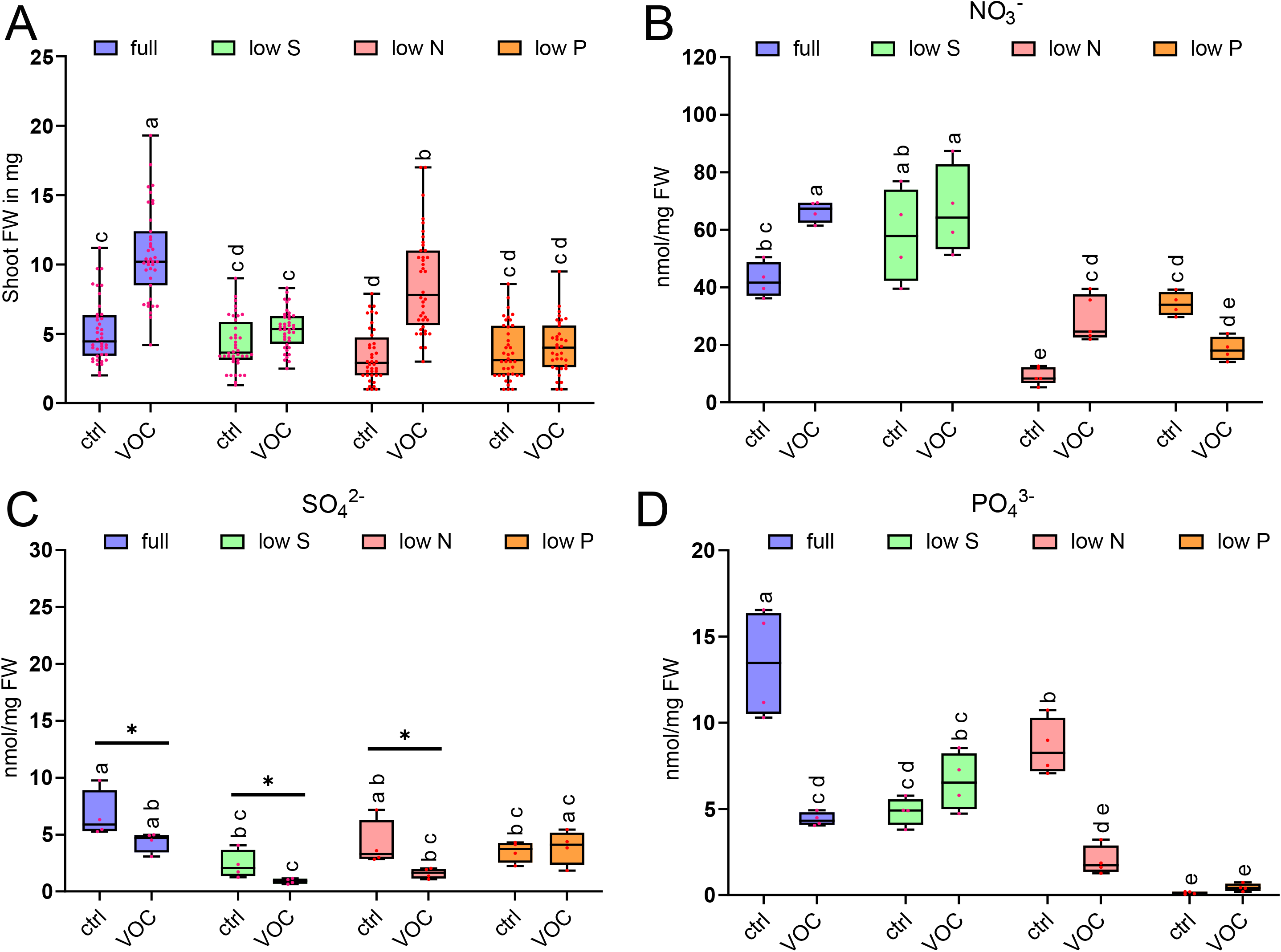
Effect of nutrient limitation on fresh weight and shoot anion levels upon 10-day 16SC VOC exposure. **A)** Fresh weight (mg) of *Arabidopsis thaliana* seedlings following 10 days of 16SC VOC treatment under sulfur, nitrate, and phosphorus deficiencies. Letters indicate statistically significant differences between group means (two-way ANOVA and Tukey’s post-hoc test, *p* < 0.05, media: F(1, 306) = 129.6, p < 0.001, treatment: F(3, 306) = 42.46, p < 0.001, interaction: F(3, 306) = 25.40, p < 0.001, n = 40; 4 independent replicates with 10 seedlings each). Shoot anion levels of 10-day 16SC VOC-treated plants measured for **B)** nitrate, **C)** sulfate, and **D)** phosphate levels. Letters indicate statistically significant differences between means (two-way ANOVA and Tukey’s post-hoc test, *p* < 0.05, B) media: F(3, 26) = 48.00, p < 0.001, treatment: F(1, 26) = 9.00, p = 0.006; interaction: F(3, 26) = 8.00, p < 0.001. C) media: (F(3, 24) = 12.04, p < 0.001), treatment: (F(1, 24) = 9.965, p = 0.004), interaction :(F(3, 24) = 1.959, p = 0.147. D) media: F(3, 24) = 47.57, p < 0.001, treatment: F(1, 24) = 43.15, p < 0.001, interaction: F(3, 24) = 25.59, p < 0.001. n = 4; each dot represents pooled anions from three plants). Asterisks (*) indicate pairwise comparison of two groups (Student’s t-test, *p* < 0.05).

### 16SC Bacterial Community VOCs Response Depends on Core Nutrient Transporters

To determine whether the nutrient effects of 16SC VOC response are due to uptake or signaling, we analysed *Arabidopsis thaliana* mutants impaired in key regulatory components and transporters for sulfate, nitrate, and phosphate. Under nutrient-sufficient conditions, growth promotion by 16SC VOCs was abolished in the mutants of the primary root sulfate transporter *sultr1;2* and the sulfur-deficiency response regulator *eil3* (Figure 2A). Similarly, growth promotion was lost was in the nitrate transporter mutant *nrt1;1* or in phosphate signaling mutants *pho2* (a negative regulator) and *phr1* (a positive master regulator) (Figure 2A). On contrary, 16SC VOCs successfully promoted growth in the absence of *nlp7*, the key regulator of the primary nitrate response. This indicates that while the VOC-mediated PGP response requires nutrient transporter machinery, it can bypass the central NLP7 signaling hub to trigger biomass increase. Ion analysis confirmed that in the genotypes where growth was promoted (WT and *nlp7*), there was a significant and specific increase in shoot nitrate levels (Figure 2B). In contrast, phosphate and sulfate levels generally decreased or remained stable, consistent with the links between nitrate accumulation and growth promotion (Figure S1A,B).

**Figure 2.**
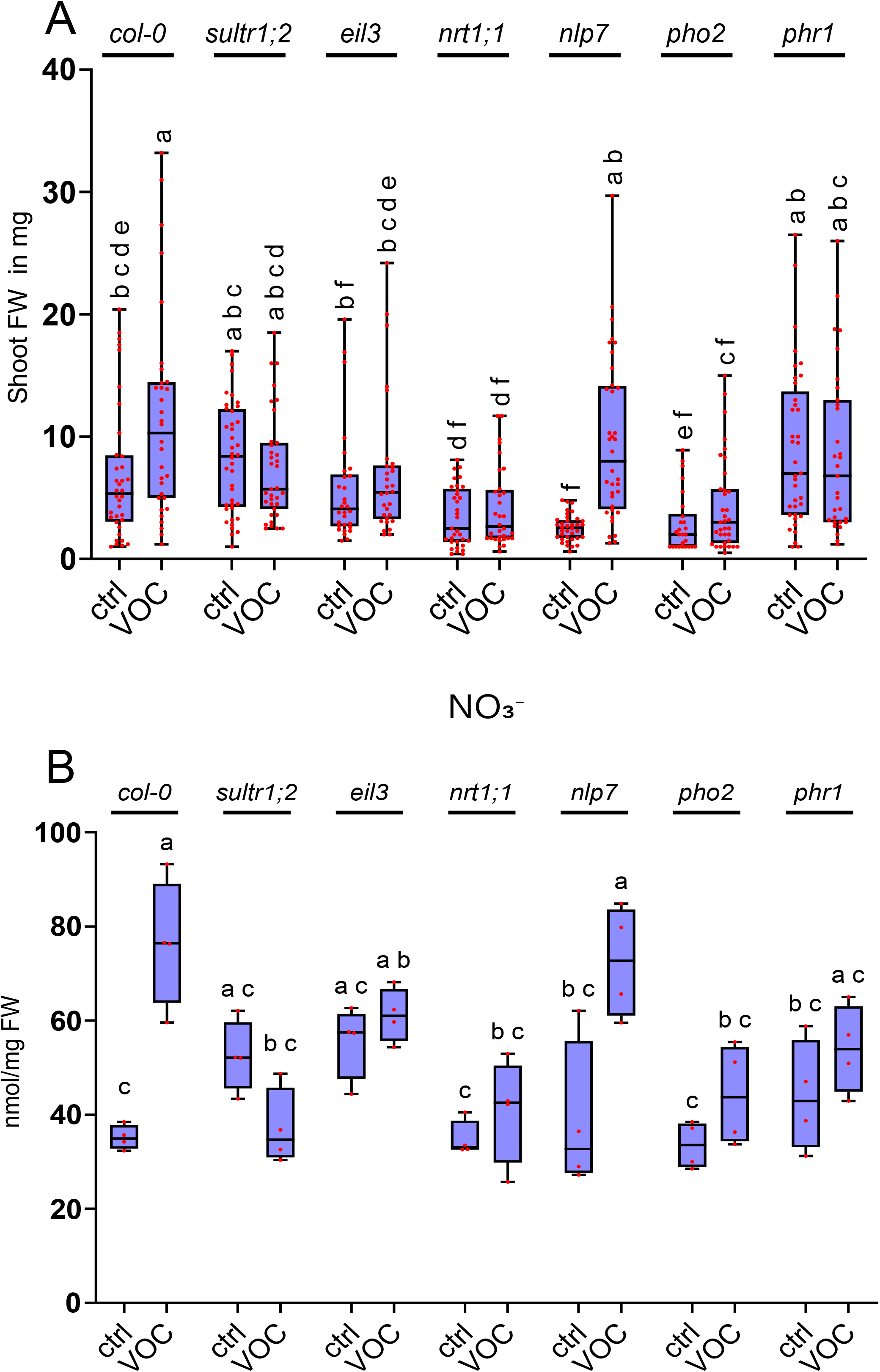
Possible involvement of nutrient pathway genes in 16SC VOC-mediated growth modulation of *Arabidopsis thaliana*. **A)** Fresh weight (mg) of *Arabidopsis thaliana* seedlings (WT, *sultr1;2, eil3, nrt1;1, nlp7, pho2,* and *phr1*) under full media conditions following 10 days of 16SC VOC treatment. Letters indicate statistically significant differences between group means (two-way ANOVA and Tukey’s post-hoc test, *p* < 0.05, genotype: F(6, 455) = 12.54, p < 0.0001; treatment: F(1, 455) = 17.57, p < 0.0001; interaction: F(6, 455) = 5.639, p < 0.0001, n = 40; 4 independent replicates with 10 seedlings each). **B)** Shoot nitrate levels of WT, *sultr1;2, eil3, nrt1;1, nlp7, pho2,* and *phr1* plants after 10 days of 16SC VOC treatment. Letters indicate statistically significant differences between means (two-way ANOVA and Tukey’s post-hoc test, *p* < 0.05, genotype: F(6, 42) = 5.921, p = 0.0002, treatment: F(1, 42) = 25.63, p < 0.0001; interaction: F(6, 42) = 7.570, p < 0.0001 n = 4; each dot represents pooled anions from three plants). Asterisks (*) indicate pairwise comparison of two groups (Student’s t-test, *p* < 0.05).

This physiological shift was mirrored by changes in gene expression (Figure 3). In WT plants, 16SC VOCs led to a significant downregulation of the sulfate transporters *SULTR1;2* and *SULTR3;5* (responsible for root-to-shoot translocation), explaining the lower sulfate accumulation in the shoot (Figure 3A, D). Interestingly, while the expression of the dual-affinity nitrate transporter *NRT1;1* remained unchanged, the high-affinity transporter *NRT2;1* was significantly downregulated (Figure 3B, E). This suggests that the high internal nitrate levels induced by VOCs may trigger a feedback mechanism to reduce high-affinity uptake. No significant changes were observed in the expression of phosphate-related genes *PHT1;1* or *PHR1* (Figure 3C, F), suggesting that the initial VOC response is primarily wired through nitrogen and sulfur metabolic reprogramming.

**Figure 3.**
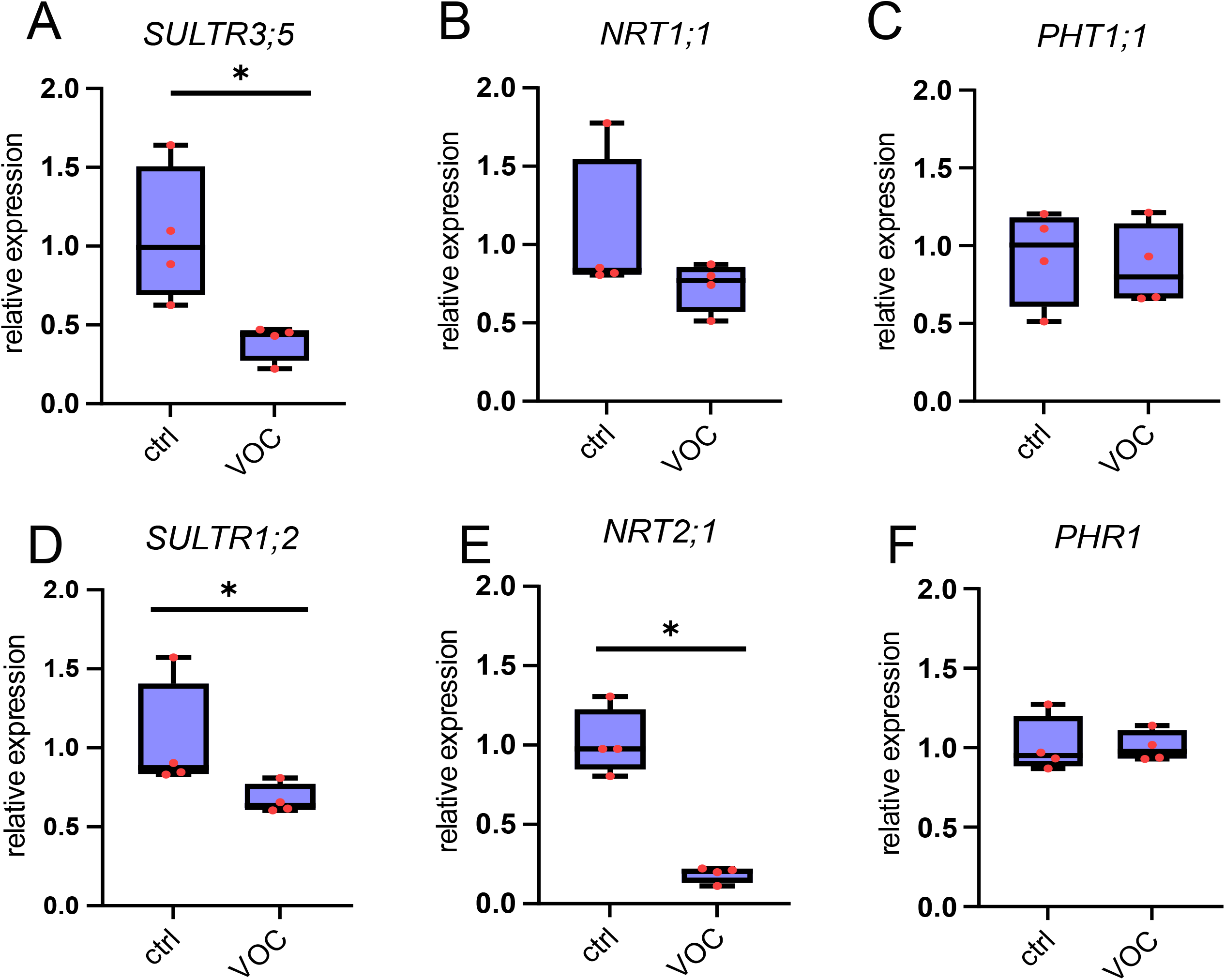
Gene expression further confirmed changes in nutrient status. Gene expression of nutrient pathway genes in *Arabidopsis thaliana* roots: **A)** *SULTR1;2*, **B)** *NRT1;1*, **C)** *PHT1;1*, **D)** *SULTR3;5*, **E)** *NRT2;1*, and **F)** *PHR1* upon 10 days of 16SC VOC treatment under full media conditions. All gene expression data were normalized to *UBIQUITIN* (*UBQ*) expression. Asterisks (*) indicate pairwise comparison of two groups (Student’s t-test, *p* < 0.05); ns = no significant difference. n = 4; each dot represents pooled gene expression from 3 plants. Reactions were performed in duplicate for four independent biological samples.

To assess whether these effects are maintained under nutrient limitation, we subjected each mutant line to a limitation of its corresponding nutrient and analyzed their growth and nutrient status (Figures S2–S4). The signalling mutants behaved in the same way in control and limitation conditions, i.e., *nlp7* gained FW upon exposure to the VOCs, while *eil3* and *phr1* did not (Figures S2A-S4A). Interestingly, while the *nrt1;1* and *sultr1;2* mutants showed a loss of growth promotion under full nutrient (Figure 2A), they regained this beneficial effect when subjected to low N or low S, respectively (Figures S3A and S2A). The response of *pho2* did not differ in deficiency from control conditions (Figures S4A). Anion levels remained relatively stable across these conditions, except the mutants in S pathway, *sultr1;2* and *eil3*, which exhibited a reduction in shoot nitrate levels upon 16SC VOC treatment (Figures S2–S4).

### Nutrient Limitation Prevents PGP Effect of 16SC Bacterial Community in Direct Bacterial Contact

The 16SC community possess a PGP effects on plants through VOCs as well as in direct contact (Türksoy et al., 2025; Wippel et al., 2021). To find out how nutrient limitation affects the direct 16SC growth promotion, we employed the same nutrient limitations while inoculating the 16SC directly into the media. After 10 days of DBC, growth promotion was lost under all three nutrient-limited conditions (Figure 4A). Strikingly, the nutrient status of the plant shoots remained stable and did not change significantly upon 16SC DBC inoculation, although the respective deficiencies were clearly reflected in the absolute shoot anion levels (Figure S5A,B,C)

**Figure 4.**
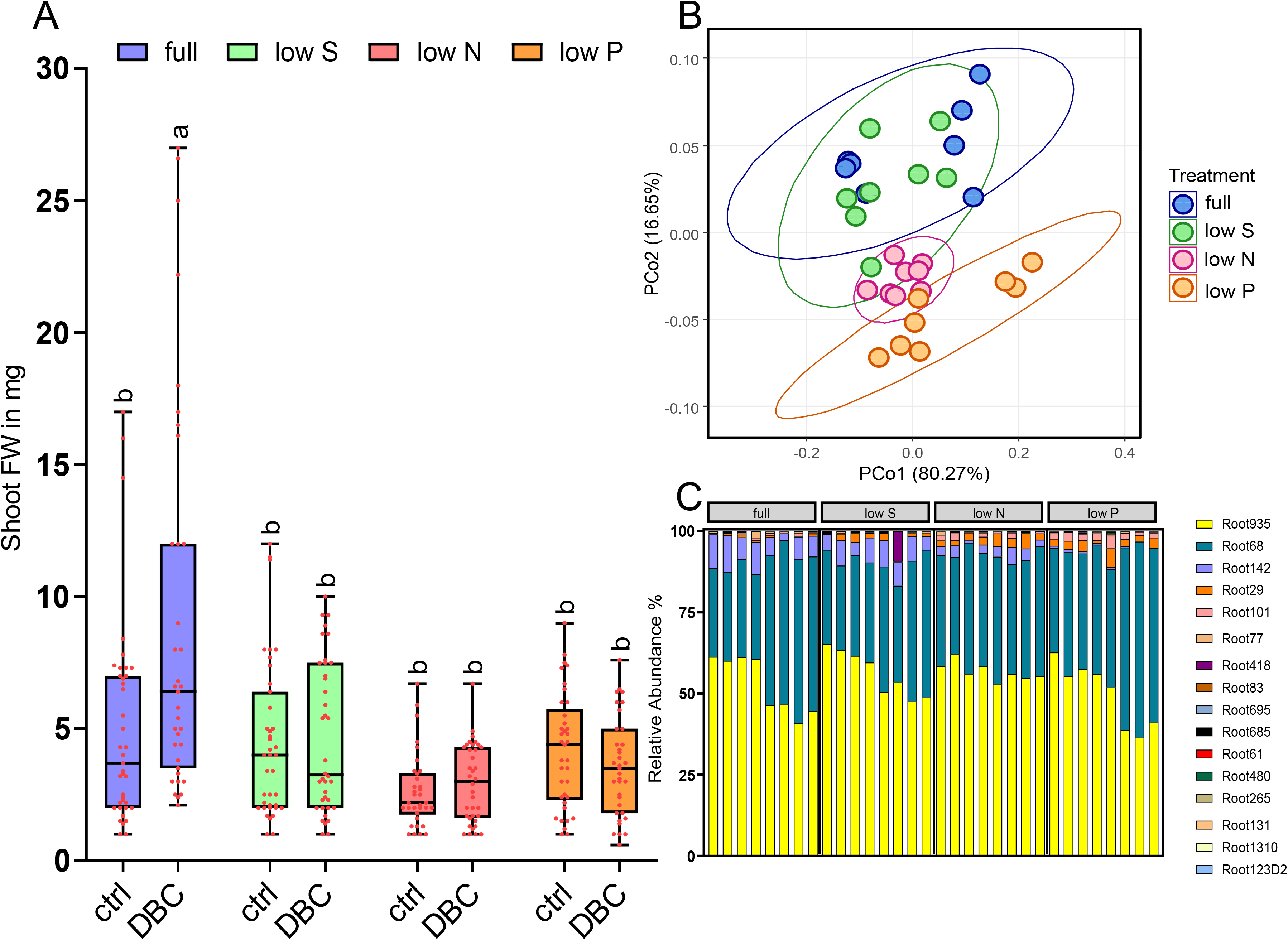
Effect of nutrient limitation on fresh weight and community composition upon 10-day 16SC DBC exposure. **A)** Fresh weight (mg) of *Arabidopsis thaliana* seedlings under sulfur, nitrate, and phosphorus deficiencies following 10 days of 16SC DBC treatment. Letters indicate statistically significant differences between means (two-way ANOVA and Tukey’s post-hoc test, *p* < 0.05, media: F(3, 285) = 19.46, p < 0.0001; treatment: F(1, 285) = 5.807, p = 0.0166; interaction: F(3, 285) = 6.861, p = 0.0002, n = 40; 4 independent replicates with 10 seedlings each). **B)** Principal coordinates analysis (PCoA) of Bray–Curtis dissimilarities showing the composition of a 16-member bacterial synthetic community (SynCom) colonizing the roots of *Arabidopsis thaliana*, grown on agar media under full nutrient conditions, low sulfur, low nitrate, and low phosphate for 10 days. **C)** Bar plot showing the relative abundance of individual strains within the 16-member SynCom colonizing *Arabidopsis thaliana* roots under full nutrient, low sulfur, low nitrate, and low phosphate conditions after 10 days of co-cultivation on an agar plate system. Relative abundances were determined by 16S rRNA profiling. Each condition includes 8 biological replicates.

To reveal whether nutrient limitations primarily disrupt bacterial colonization and community assembly or interfere with the functional growth-promoting effects of the colonized strains, we performed 16S rRNA amplicon sequencing. Interestingly, based on PcoA analysis, while low S treatment resulted in a bacterial root colonization profile similar to the 16SC full media, both low N and low P conditions produced distinct community profiles (Figure 4B).

The 16S rRNA profiling revealed that the SynCom architecture is highly resilient to nutrient limitation, dominated across all conditions by two major strains, R68 (*Pseudomonadaceae*) and R935 (*Flavobacteriaceae*), which maintained their dominance regardless of the nutrient status (Figure 4C, Figure S6A, B). Instead, the most significant shifts occurred within the minor strains (Figure 4C) across the treatments. As expected, the most extensive changes in root colonization were observed under low P, characterized by a decrease in R142 (*Rhizobiaceae*) (Figure S6C) and a significant increase in the abundance of several other strains, including R29 (*Comamonadaceae*), R101 (*Intrasporangiaceae*), R695 (*Phyllobacteriaceae,* R265 (*Mycobacteriaceae*), and R685 (*Hyphomicrobiaceae*) (Figure S6). Although low S had a minimal overall impact on colonization compared to the full media, it specifically led to reduced colonization of R101 (*Intrasporangiaceae*). Furthermore, low N treatment significantly increased the relative abundance of R29 (*Comamonadaceae*) (Figure S6D). Taken together, these findings demonstrate that limitations in each of the three major macro-nutrients reshape the root microbiome composition in a distinct manner. Crucially, the lack of significant structural disruption under low S suggests that the observed loss of growth promotion is likely driven by a nutrient deficiency-induced shift in bacterial functionality rather than altered colonization. Conversely, the extensive, nutrient-specific recruitment of minor strains under low P was ultimately insufficient to compensate for this functional collapse, failing to rescue the plant growth phenotype.

### DBC-Mediated Growth Promotion is Dependent on Nutrient Master Regulators

To dissect how the primary transporters and regulators of S, N, and P homeostasis influence the beneficial effects of 16SC during DBC, we evaluated the same set of mutants previously tested under VOC treatment. This comparison aimed to determine whether nutritional status interacts differently with long-distance (volatile) and short-distance (contact-dependent) microbial signals. Interestingly, we observed distinct differences in the growth responses of specific mutants to 16SC DBC compared to VOC exposure. In contrast to the VOC treatment, *sultr1;2* and *nrt1;1* exhibited significant growth promotion under DBC conditions. On contrary, the regulatory mutants *eil3* and *nlp7* showed a loss of growth promotion (Figure 5A). This suggests that while growth promotion via VOCs may rely more on transporter-mediated mechanisms, the plant’s response to direct physical interaction is more dependent on core regulatory genes. Conversely, both P signaling mutants, *phr1* and *pho2,* failed to show a growth-promotion effect under DBC, a result consistent with our observations under VOC treatment (Figure 5A). This indicates that the P signaling pathway is a crucial and conserved hub for both DBC- and VOC-mediated interactions with 16SC. Furthermore, as previously observed in the wild-type (WT) (Figure S5A,B,C), no significant changes were detected in shoot anion levels in the WT or in any of the transporter and regulator mutants under DBC conditions (Figure 5B and Figure S7A, B).

**Figure 5.**
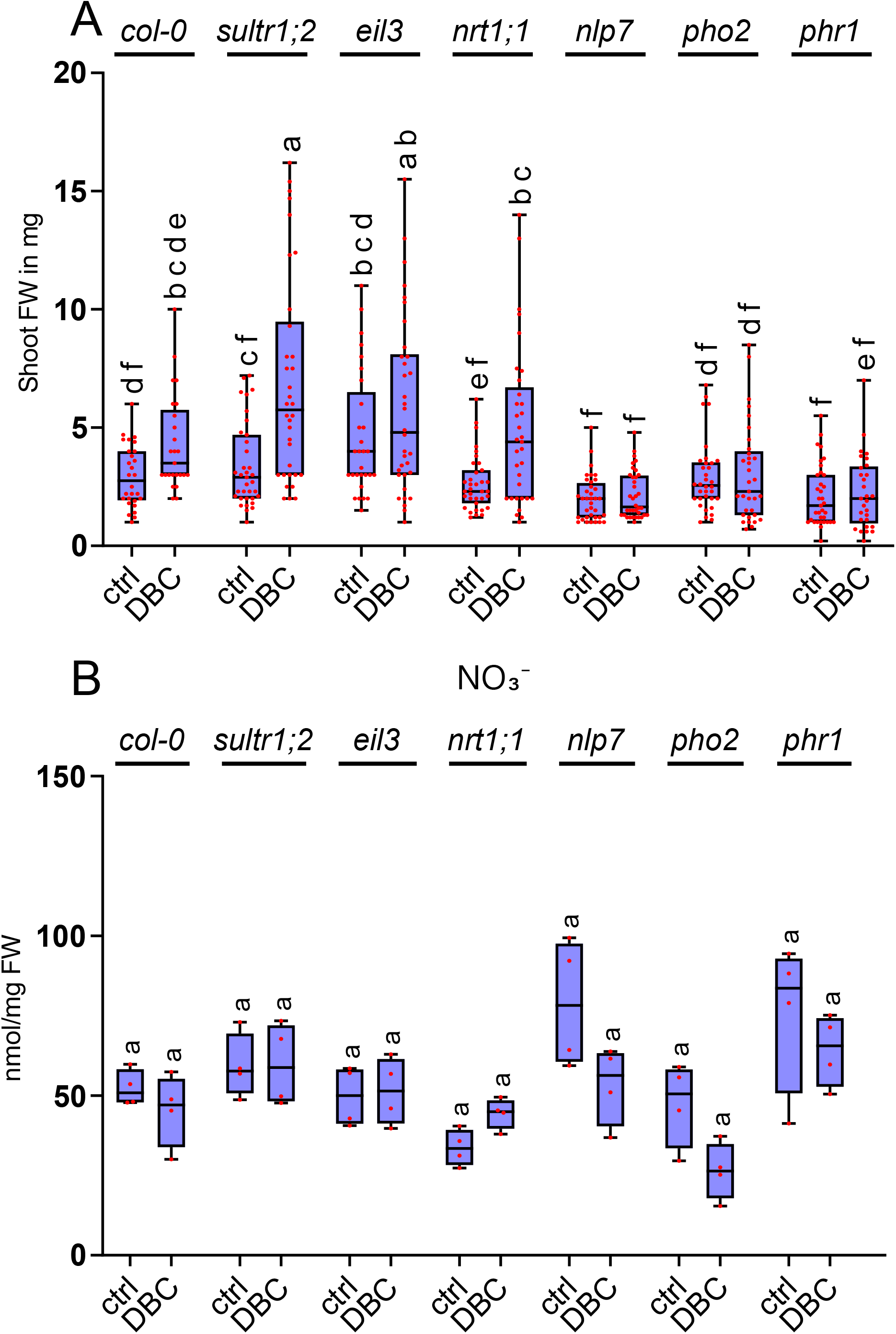
Possible involvement of nutrient pathway genes in 16SC DBC-mediated growth modulation of *Arabidopsis thaliana*. **A)** Fresh weight (mg) of *Arabidopsis thaliana* seedlings (WT, *sultr1;2, eil3, nrt1;1, nlp7, pho2,* and *phr1*) under full media conditions following 10 days of 16SC DBC treatment. Letters indicate statistically significant differences between means (two-way ANOVA and Tukey’s post-hoc test, *p* < 0.05, genotype: F(6, 442) = 20.58, p < 0.001, treatment: F(1, 442) = 33.93, p < 0.001, interaction: F(6, 442) = 5.359, p < 0.001, n = 40). **B)** Shoot nitrate levels of WT, *sultr1;2, eil3, nrt1;1, nlp7, pho2,* and *phr1* plants after 10 days of 16SC DBC treatment. Letters indicate statistically significant differences between means (two-way ANOVA and Tukey’s post-hoc test, *p* < 0.05, genotype: F(6, 42) = 8.377, p < 0.0001, treatment: F(1, 42) = 5.031, p = 0.0302, interaction: F(6, 42) = 2.139, p = 0.0687, n = 4; each dot represents pooled anions from three plants).

### Metabolic Reprogramming and Differential Exudation Patterns Distinguish Volatile-Mediated from Contact-Dependent Interactions

To elucidate how the host plant reprograms its metabolism in response to microbial cues under diverse nutrient limitations, we conducted metabolite profiling of the shoots, roots, and exudates under both 16SC VOC and DBC treatments. Under 16SC VOC exposure, the metabolic profiles of the three analyzed compartments separated sharply from their respective mock controls in all nutrient conditions (Figure S8). Comparing the effects of nutrient deficiencies, the shoot, root, and exudate metabolomes under low P exhibited the most pronounced separation from all other treatment groups (Figure S8). Interestingly, growth-promoting conditions (the full media control and low N) under VOC exposure displayed convergent shoot metabolic profiles that clustered tightly together, remaining distinctly segregated from the non-growth promoting low S and low P conditions (Figure S8A). In the mock controls, the overall metabolite profiles were not largely affected by limitations (Figure S8). However, low S was characterized by a specific decline in S-containing amino acids (methionine and cysteine) in shoots, while low N triggered a predictable reduction in N-containing amino acids and their derivatives, such as L-valine, glutamine, asparagine, and beta-alanine (Figure S9A).

Under VOC treatment, shoots of plants grown under full media and low N growth promoting conditions shared highly similar profiles for amino acids, derivatives, and TCA cycle intermediates (Figure S9A). Conversely, non-growth promoting low S and low P limitations under VOC exposure exhibited a distinct metabolic signature marked by a robust coordinated increase in amino acids compared to full media and low N counterparts (Figure S9A).

In the root compartment, metabolic reconfiguration was more tightly coupled to specific nutrient deficiencies (Figure S8B), with the most dramatic shifts being captured under VOC-treated, P-limited conditions (Figure S8B). Specifically, low P roots under VOC exposure accumulated high levels of amino acids, derivatives, pipecolic acid, and shikimic acid, while simultaneously displaying a substantial depletion of sterols, scopoletin, and salicylic acid (Figure S9B). While the baseline exudation profiles of mock plants did not vary drastically across nutrient deficiencies (Figure S8C), the introduction of VOCs induced a striking inversion of metabolite trends compared to their respective mock controls (Figure S8C). Specifically, VOC perception drove a generalized accumulation of amino acids in the root exudates, accompanied by a decline in TCA cycle intermediates and sugars,except under low P, where the distinct metabolic behavior of the system was most pronounced (Figure S9C). This suggests that while VOCs fundamentally prime the plant to enrich its rhizosphere with amino acid-rich substrates, specific metabolic adjustments are concurrently deployed to mitigate nutrient-specific stress.

In contrast to the VOC-mediated responses, 16SC DBC triggered rigid, compartment-specific metabolic trajectories. In the shoots, the metabolic profiles for each nutrient deficiency separated clearly from one another and from their mock controls (Figure S8D). In the roots, low N induced the most prominent metabolic divergence (Figure S8E). Crucially, while mock exudates clustered tightly regardless of nutrient status, DBC inoculation fundamentally reconfigured the exuded metabolome, with low P conditions forming a highly distinct cluster (Figure S8F).

At the individual tissue level, DBC exposure altered shoot profiles in a media-dependent manner (Figure S10A). Low S and low P led to a marked accumulation of amino acids and derivatives relative to their own controls and the DBC-treated full or low-N media. While low N shoots specifically accumulated sugars, the most extensive metabolic restructuring occurred under low P DBC conditions, which triggered an overall upregulation of the TCA cycle, sterols, vitamins, shikimic acid, and various other pathways (Figure S10A).

In the root tissue, low N DBC samples segregated completely from the remaining media profiles (Figure S8E). Under DBC, low N led to a significant decrease in multiple N-containing amino acids, while low S DBC specifically depleted S-containing amino acids relative to their mock controls (Figure S10B).

In the exudates, mock treatment separated from DBC treated condition sharply under all deficiency condition (Figure S10F). Under low S DBC conditions, nearly all sugars and amino acid derivatives in the exudates dropped significantly below mock levels (Figure S10C). Conversely, low P DBC conditions induced a massive exudation spike, significantly increasing the levels of amino acids and derivatives (e.g., glycine, gamma-aminobutyric acid, 2-aminobutyric acid), TCA cycle intermediates (citric acid, oxoglutaric acid, pyruvic acid), sugars (D-ribose, L-rhamnose), phenylpropanoids, pipecolic acid, and jasmonic acid (Figure S10C). Strikingly, this exudate remodeling under DBC correlates tightly with our 16S bacterial community profiling (Figure 4B, C), where low P drove the most extensive restructuring of minor strains, followed by low N, while low S maintained a structural assembly similar to the full media while modulating a single minor strain (Figure 4B, C). These findings suggest that while host-mediated exudation plays a central role in driving community assembly, these reconfigurations under severe nutrient stress are insufficient to sustain or recruit the specific functional partners required to rescue the growth phenotype.

A direct comparison of specific metabolites revealed a systemic, inverse coordination between the 16SC VOC and DBC treatments across all plant compartments and exudates (Figure 6A, B,C). In the shoots, VOC exposure induced a stronger accumulation of the majority of amino acids and derivatives compared to full media, with L-citrulline and L-histidine reaching peak abundances under VOC relative to DBC (Figure 6A). Conversely, TCA cycle intermediates were systematically lower under VOC exposure across all media conditions compared to DBC (Figure 6A).

**Figure 6.**
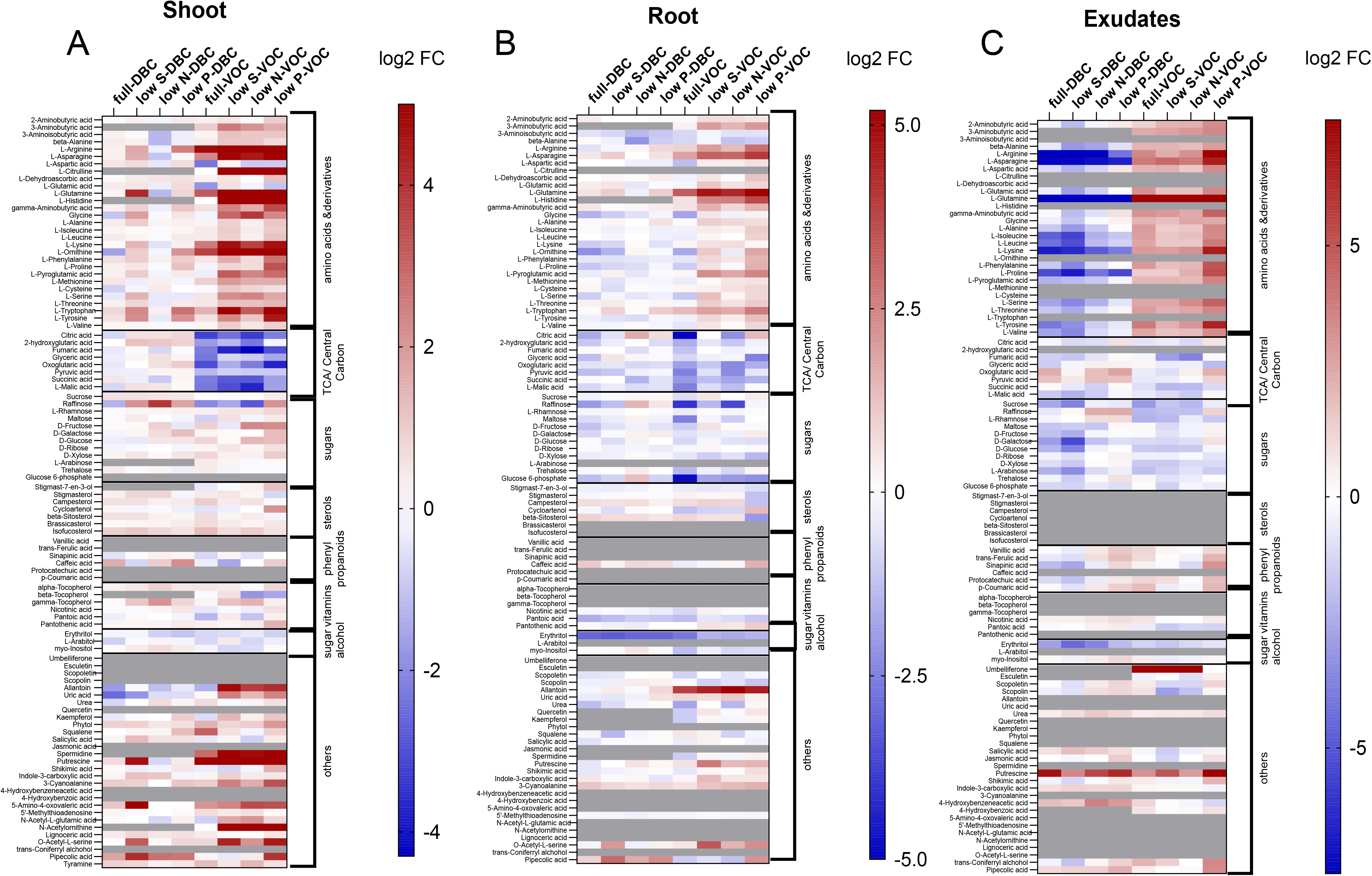
Plant compartments and exudate metabolite difference between VOC and DBC treatment under deficiency. The heatmaps show the log_2_ fold change between VOC or DBC-treated and mock-treated plants under their respective media condition.

A parallel trend was captured in the roots, where amino acids and derivatives, most notably 3-aminobutyric acid, L-arginine, L-glutamine, glycine, and L-pyroglutamic acid, accumulated highly under nutrient-limited VOC treatments compared to both full media VOC and DBC setups. Furthermore, allantoin levels were exceptionally higher in the roots under all VOC conditions compared to DBC (Figure 6B).

This opposite behavior was highly visible in the root exudates, which manifested a clear footprint of putative microbial consumption of exuded metabolies versus volatile priming. While amino acids accumulated significantly during VOC perception, they were severely depleted upon direct bacterial interaction (Figure 6C). This phenomenon was best exemplified by L-glutamine, which spiked under all VOC treatments but was sharply exhausted in all DBC conditions. Similarly, L-arginine concentration dropped dramatically under DBC relative to VOC, with the sole exception of the low P DBC treatment. Additionally, VOCs specifically enhanced the exudation of umbelliferone across all treatments except under low P (Figure 6C).

Taken together, the systematic depletion of specific metabolites under DBC strongly implies that colonizing bacteria actively utilize these plant-derived substrates during physical interaction in the rhizosphere. Conversely, the accumulation captured in the VOC exudate profiles supports two non-exclusive biological scenarios: either the physical isolation of the SynCom in the VOC setup prevents the biological drawdown of these metabolites, allowing them to accumulate, or VOCs function as an airborne systemic signal that actively prompts the plant to accelerate the exudation of specific recruiting agents. This pre-emptive exudation strategy suggests an evolutionary mechanism whereby the host plant, upon sensing volatile microbial signals at a distance, structurally enriches the rhizosphere to attract or sustain beneficial partners ahead of physical contact.

## Discussion

### Bacterial Community Volatiles Modulate *Arabidopsis thaliana* Growth Under Nutrient Limitation via Specific Genetic and Metabolic Checkpoints

As previously shown in Türksoy et al. (2025) (Türksoy et al., 2025), 16SC community promoted plant growth through community-emitted VOC blend. DMDS was identified as the dominant volatile in this blend, together with several additional sulfur-containing, nitrogen-containing, and terpene-derived compounds. Since these compounds could represent alternative sources of these mineral nutrients, as shown by Meldau et al., (2013) (Meldau et al., 2013) for DMDS produced by *Bacillus* strain B55, the interaction of VOC treatment with nutrient pathways is of great interest. Indeed, for instance, sulfur has increasingly been recognized as an important factor in plant–microbe interactions in dual role as signal and nutrient (Elkatmis et al., 2026; Mukherjee, Han, et al., 2025). Moreover, previous studies showed that *Bacillus* strain GB03 can increase rhizosphere acidification, enhance iron availability, and affect iron uptake mechanisms in plants (del Carmen Orozco-Mosqueda et al., 2013; H. Zhang et al., 2009). Based on these findings, we investigated whether the beneficial effects of the 16SC VOC blend are dependent on the host’s nutritional context, using low N, low P and low S limitations as distinct environmental regimes

Interestingly, although DMDS was identified as the dominant volatile emitted by the 16SC (Türksoy et al., 2025), growth promotion was not observed under sulfur deficiency, whereas a positive effect was detected under low nitrogen conditions (Figure 1A). This may indicate that the concentration or duration of DMDS exposure was insufficient to replenish the plant sulfur pools. Alternatively, the VOC blend may trigger broader reprogramming of nutritional pathways rather than directly supplying sulfur. Supporting this interpretation, Türksoy et al., (2025) (Türksoy et al., 2025) also showed that *oastlA* and *oastlBC* mutant plants, which cannot assimilate DMDS, still displayed growth promotion when exposed to the community VOC blend.

One of the most striking observations was that plants exposed to the 16SC VOCs still displayed growth promotion under nitrate limitation, together with similar nitrate, sulfate, and phosphate levels in shoots compared to full nutrient condition (Figure 1). This suggests that VOCs may either serve as alternative nitrogen sources or induce pathways that improve nitrogen use efficiency. Previous studies have shown that bacterial community composition changes under N stress and is correlated with plant root metabolites. Additionally, bacterial community richness has been positively linked to nitrogen use efficiency in sorghum, and under N stress, *Sphingopyxis* has been shown to enhance shoot biomass and nitrogen acquisition (Chai et al., 2024; Li et al., 2026). Moreover, Chen et al., (2024) (Chen et al., 2024) showed that *Bacillus velezensis* VOCs promote growth and enhance nitrogen uptake from the environment by increasing nitrate uptake in *Arabidopsis thaliana* and rice, consistent with our findings. These observations highlight the potential of bacterial VOCs for reducing nitrate-based fertilizer use, which is associated with major environmental problems (Guo et al., 2010; Liu et al., 2013).

Further analysis using mutant lines under both full and nutrient-limited conditions demonstrated that nutrient transporters play important roles in VOC-mediated growth promotion. Under full nutrient conditions, the *nrt1;1* and *sultr1;2* mutants lost the growth-promoting effect (Figure 2A), whereas under their respective nutrient limitations, growth promotion was restored (Figures S2A, S3A). Interestingly, the transcript levels of these transporters were not affected by the VOC treatment (Figure 3). This suggests that plants employ different transporters or nutrient acquisition pathways under varied nutrient availability. *NRT1;1* has previously been implicated in plant–microbe interactions associated with nitrogen use efficiency (Zhang et al., 2019) ^39^, but much les is known of its function in the context of VOC-mediated growth modulation. Interestingly, Chen et al., (2024) (Chen et al., 2024) showed that *B. velezensis* induced *NRT2;1* transcript levels and the PGP effect was dependent on NLP7, while 16SC suppressed *NRT2;1* expression and *nlp7* mutant increased growth upon 16SC VOC treatment (Figure 2A, 3E). It seems that different VOCs trigger different signalling pathways and employ different transcriptional networks to converge on nitrate accumulation. This difference may also reflect the complexity of community-emitted VOC mixtures versus VOCs from single strains.

A strong correlation was observed between growth promotion under full media and low nitrogen conditions, particularly in the *nlp7* mutant, where growth promotion was consistently associated with increased shoot nitrate accumulation (Figure 2A, B). This indicates that nitrate accumulation may contribute to the observed growth-promoting activity. Interestingly, plants bypassed the major nitrate regulator *NLP7*, suggesting that pathways downstream of nitrate uptake or signaling through *NRT1;1* may be involved. However, this relationship was not observed under low sulfur and low nitrogen conditions in *nrt1;1*, *sultr1;2*, and *nlp7* mutants, where growth promotion still occurred without significant changes in shoot nitrate levels (Figure S2A, B and S3A, B). These findings imply that additional transporters and nutrient signaling pathways may become activated under nutrient limitation to sustain VOC-mediated growth promotion. In contrast, regulatory mutants such as *eil3*, *pho2*, and *phr1*, which fail to maintain growth promotion under nutrient limitation, may represent critical regulatory nodes integrating nutrient status with VOC-mediated growth responses.

Metabolite profiling further supported these observations. Growth-promoting conditions under full media and low nitrogen clustered closely together, whereas non-growth-promoting conditions under low sulfur and low phosphate formed distinct metabolic profiles, particularly in amino acids and their derivatives in shoots (Figure S8 and S9A). Similar nitrate accumulation patterns suggest that under low nitrate conditions plants can effectively reprogram metabolism and improve nutrient use efficiency, whereas sulfur and phosphate limitation disrupt these processes. The increased accumulation of amino acids and related compounds under low sulfur and phosphate conditions may interfere with VOC-mediated nutrient assimilation. Interestingly, under low phosphate conditions, VOC treatment increased pipecolic acid and jasmonic acid levels, metabolites associated with pathogen responses and activation of plant immunity. This regulation was specific for low P conditions; under all other conditions the levels of pipecolic acid and related metabolites decreased following VOC treatment, consistent with our previous study showing reduction in camalexin content upon VOC exposure (Türksoy et al.,2025) ^26^. Notably, P limitation increased the accumulation of these metabolites even in the mock control. Indeed, root microbiota regulate both plant immunity and phosphate stress responses; therefore, these findings may indicate that VOCs are part of this regulation under low P (Castrillo et al., 2017).

### VOC- and DBC-Mediated 16SC Interactions Affect Core Nutrient Pathways Differently

Loss of growth promotion under all nutrient deficiency conditions during DBC interaction with the 16SC was surprising (Figure 4A). Finkel et al., (2019) (Finkel et al., 2019) previously showed that phosphate limitation reduces shoot size while reshaping the plant-associated microbiome through phosphate starvation responses. In contrast, (Mukherjee, et al., 2025) ^42^ demonstrated that microbiota promoted plant growth under sulfur deficiency through glutathione supply. Unlike these findings, our 16SC did not confer growth promotion under sulfur deficiency during DBC interaction. Differences between experimental systems may partly explain these discrepancies. Whereas (Mukherjee, et al., 2025) ^42^ used a vermiculite-based system, our experiments were conducted on plates, which may delay or limit observable growth-promoting effects. Moreover, our experimental system employed mild S limitation, whereas the growth promotion was achieved under more severe S deficiency conditions.

Interestingly, in contrast to VOC-mediated interactions, *nrt1;1* and *sultr1;2* mutants still benefited from DBC (Figure 5A). Additionally, anion levels, particularly nitrate levels, remained stable during DBC interaction, unlike the nitrate accumulation observed during VOC-mediated growth promotion (Figure 5B). This contrasts with (Zhang et al., 2019), who associated rice nitrogen use efficiency with *NRT1;1B*-mediated root microbiome assembly and showed that ammonification processes were reduced in *nrt1;1* mutant plants. Together, these results suggest that plant regulatory mechanisms involved in sensing and responding to bacterial colonization may play a larger role during DBC interaction than direct nutrient provision or enhanced nutrient uptake.

Loss of growth promotion under phosphate limitation and in *pho2* and *phr1* mutants during both VOC and DBC interactions demonstrates that phosphate signaling is essential for both communication modes. In contrast, the differential responses observed for nitrogen- and sulfur-related signaling pathways indicate that these pathways contribute in a communication-type-specific manner. These findings align with previous studies showing that phosphate starvation responses strongly influence microbiome assembly (Finkel et al., 2019).

Metabolite profiling of shoots, roots, and exudates further demonstrated that plants extensively reprogram metabolism under stress conditions, likely to recruit beneficial bacteria (Figure S10A–C). Plants are known to release metabolites as a “cry for help” strategy to maintain fitness under stress and pathogen attack (Rizaludin et al., 2021). This phenomenon was clearly reflected in both the exudate profiles and bacterial community shifts observed during 16SC-DBC interaction (Figure 4B,C; Figure S10C). Phosphate limitation induced the strongest changes in both exudate composition and bacterial community structure, in agreement with previous results (Finkel et al., 2019). Nevertheless, despite these metabolic and microbial shifts, plants still failed to exhibit growth promotion under nutrient stress. This may indicate that the experimental duration was insufficient to observe beneficial effects which may be observable under soil conditions following long-term exposure.

### VOC versus DBC Metabolic Differentiation

It is well established that plants secrete metabolites that act as signals and nutrient sources to facilitate colonization (Hu et al., 2018; Jacoby et al., 2021). Amino acids and their derivatives play particularly important roles during early bacterial colonization through recognition by bacterial chemoreceptors (Allard-Massicotte et al., 2016; Moormann et al., 2022; Yang et al., 2015). Accordingly, amino acids were depleted during DBC interaction but accumulated during VOC treatment, suggesting fundamentally different plant responses to these two communication modes (Figure 6C). VOC exposure appeared to stimulate the secretion and accumulation of metabolites that may facilitate future colonization, whereas direct bacterial presence resulted in depletion of these compounds. Notably, glutamine levels in exudates drastically decreased during DBC interaction but strongly increased during VOC treatment (Figure 6C). This observation is consistent with recent findings by Tsai et al., (2025) (Tsai et al., 2025), who showed that localized glutamine levels shape spatial root microbial colonization. Similarly, VOCs emitted by a single bacterial strain promoted rice growth by 83% while altering plant metabolism, particularly amino acid metabolism (Almeida et al., 2023). Specifically, the levels of arginine, asparagine, and glutamine increased, suggesting a potential effect on nitrogen assimilation.

Furthermore, several studies demonstrated that specific amino acids and metabolism are important for the growth-promoting function of *Pseudomonas* strains in *Arabidopsis thaliana*, which also appeared as dominant taxa in our community profiling (Cheng et al., 2017; Cole et al., 2017). Likewise, Koprivova et al., (2025) (Koprivova et al., 2025) showed that exudates from *Arabidopsis thaliana* treated with the PGP *Pseudomonas* strain CH267 exhibited a significant decrease in amino acids. This may indicate a robust and conserved pathway influencing bacterial colonization, both at the level of single strains and within microbial communities. Moreover, root and exudate sugar levels decreased under both experimental settings, which may further suggest microbial consumption of specific metabolites prior to and during colonization, in both single-bacterium and community-mediated interactions. However, it cannot be excluded that these differences result from different regulation of exudation dynamics by DBC and VOC treatment. Under DBC conditions, defense-associated metabolites such as pipecolic acid and salicylic acid accumulated, suggesting activation of immune-related responses. In contrast, VOC exposure did not trigger these responses (except low P), potentially reflecting a priming state that prepares plants for bacterial colonization without activating full defense pathways, as previously suggested (Türksoy et al., 2025).

Overall, this study demonstrates that nutrient status affects VOC-mediated and direct bacterial communication differently. Differences in growth promotion, ion accumulation, bacterial community composition, and especially amino acid profiles in exudates reveal distinct regulatory mechanisms. While previous studies mainly focused on DBC-induced changes in exudation, our findings highlight VOCs as powerful tools capable of reprogramming plant metabolism, shaping microbiome assembly, and improving stress tolerance. In particular, the strong growth-promoting effect observed under low nitrogen conditions suggests that VOC-based approaches could potentially be engineered for use in nitrate-depleted soils. However, further studies are required to fully understand the underlying mechanisms.

## Material & Methods

### Plant Material, Growth Media, and Growth Conditions

*Arabidopsis thaliana* ecotype Columbia-0 (Col-0) was used as the wild type (WT) in all experiments. Six mutants defective in nutrient-related genes were used: *sultr1;2* (Yoshimoto et al., 2002), *eil3* (SALK_089129) ^28^, *nrt1;1* (Ho et al., 2009; Ristova & Kopriva, 2022), *nlp7* (SALK_114886) ^31^*, phr1* (Bustos et al., 2010), and *pho2* (Aung et al., 2006). Plant material was grown on plates with modified Long Ashton medium solidified with agarose under full nutrient supply (Control), low sulfate (low S; 0.015 mM sulfate, see Table S1, low nitrate (low N; 1 mM nitrate), or low phosphate (low P; 0.06 mM). For stratification and germination, the medium was supplemented with 0.5% (w/v) sucrose. Seeds were sterilized using chlorine gas before sowing onto the corresponding media in square plates, which were sealed with 3M Micropore tape. Plates were stratified at 4°C for 3 days in darkness and then transferred to a growth chamber (16 h light/8 h dark cycle, 21°C/19°C, PPFD 100 μmol m⁻² s⁻¹) for 5 days of germination.

### Bacterial Material, Growth Media, and Growth Conditions

Sixteen bacterial strains representing different families of the *Arabidopsis thaliana* root microbiome were used in this study. This 16SC was derived from SynCom At-SC3 (Wippel et al., 2021), with minor modifications: Root1221 was replaced by Root29, and Root1485 was replaced by Root418 (Türksoy et al., 2025) (Table S2). Each strain was grown for 2–5 days on 50% Tryptic Soy Agar (TSA) plates at 28°C prior to plant inoculation. One day before inoculation, liquid cultures were prepared by transferring several colonies with a sterile pipette tip into 5 mL of 50% Tryptic Soy Broth (TSB) in culture tubes. Cultures were incubated at 28°C with shaking at 200 rpm.

### Co-cultivation of Plants and Bacteria

#### VOC Exposure

Combi-plates for VOC exposure without direct contact between plants and bacteria were prepared as previously described (Türksoy et al., 2025). Fifty mL of control or nutrient-limited media was poured into square Petri dishes and a small Petri dish containing 3 mL of molten 50% TSA was attached on the lower half of the plate. Bacterial cultures were centrifuged for 10 min at 2500 rpm and 22°C, the pellets were resuspended in 1 mL of 0.9% (w/v) NaCl solution, and adjusted to OD600 = 1. Equal volumes of each strain were combined to generate the 16SC. For VOC treatments, 100 μL of 0.9% NaCl (control) or 100 μL of 16SC suspension was spread onto the 50% TSA agar in the small Petri dish and allowed to dry. Ten seedlings were transferred to the upper part of the square Petri dishes. Plates were sealed with 3M Micropore tape, positioned vertically, randomized, and incubated in a growth chamber for 10 days (16 h light at 21°C / 8 h dark at 19°C, PPFD 100 μmol m⁻² s⁻¹).

#### Direct Bacterial Contact

For direct contact treatments, strains were adjusted to OD600 = 0.2 using 0.9% NaCl. Then, 150 μL of bacterial suspension was added to 50 mL of 40°C warm control or nutrient-limited agar media to obtain a final density of OD600 = 0.0005. The mixture was poured into square Petri dishes and stored at room temperature overnight before seedling transfer. Control plates received 150 μL of 0.9% NaCl instead of bacterial suspension. Ten seedlings were transferred to the upper part of the square Petri dishes. Plates were sealed with 3M Micropore tape, positioned vertically, randomized, and incubated in a growth chamber for 10 days.

### Anion Analysis

For nitrate, phosphate, and sulfate quantification, 1 mL of sterile Milli-Q water was added to 1.5 mL screw-cap tubes and placed on ice. Frozen shoot samples were homogenized in 200 μL of this water using a pestle homogenizer. Extracts were returned to the tubes and kept on ice. Samples were shaken for 1 h at 1000 rpm and 4°C, heated to 95°C for 15 min, and centrifuged at 15,000 rpm and 4°C. Samples were diluted 1:10 with sterile Milli-Q water prior to measurement. Anion concentrations were measured using a DIONEX AS-AP ion chromatography system (Thermo Scientific) equipped with a DIONEX IonPac AS22 (4 × 250 mm) column at a flow rate of 1.2 mL min⁻¹. The running buffer consisted of 4.5 mM Na₂CO₃ / 1.4 mM NaHCO₃. Data acquisition and peak identification were performed using Chromeleon™ software.

### RNA Extraction cDNA Synthesis and Quantitative PCR (qPCR)

Total RNA was isolated using standard phenol/chloroform/isoamyl alcohol extraction followed by LiCl precipitation. RNA concentration was measured using an IMPLEN NanoPhotometer N60. First-strand cDNA was synthesized from 800 ng total RNA using the QuantiTect Reverse Transcription Kit (Qiagen, Hilden, Germany). Quantitative PCR (qPCR) was performed using a BIO-RAD C1000 Touch™ Thermal Cycler with a CFX96™ Real-Time System. Gene expression levels were normalized to *UBIQUITIN* (UBQ) as a reference gene. Primer sequences are listed in Table S3. Reactions were performed in duplicates for four independent biological samples.

### Bacterial Profiling

DNA was extracted from plant roots using the FastDNA SPIN Kit for Soil (MP Biomedicals, Solon, USA) and eluted in 40 μL nuclease-free water. The communities were profiled by amplicon sequencing of the variable V5–V7 regions of the bacterial 16S rRNA gene. rRNA gene amplification, library preparation, amplicon sequencing, and raw data processing were performed by Novogene. Merged FASTQ obtained from Novogene were processed using the DADA2 (v1.38.0) package (Callahan et al., 2016) in R (v4.5.2) for quality filtering, denoising, and high-resolution identification of Amplicon Sequence Variants (ASVs). To ensure the accurate detection of the 16 specific bacterial strains used in this study, a custom ’fuzzy matching’ algorithm was implemented using the stringdist (v0.9.17) and Biostrings packages (van der Loo, 2014).

In this approach, inferred ASVs were aligned directly against a reference library containing the V5–V7 regions of the 16 target strains, which were specifically trimmed to match the sequenced amplicon length. A similarity threshold of ≥95% was applied using the Levenshtein distance metric to account for potential natural variations and sequencing bias within the V5–V7 regions, ensuring that related sequences were correctly assigned to their respective strains. Downstream data manipulation and normalization were performed using the tidyverse suite, including dplyr and tidyr. To account for variations in sequencing depth across samples, the counts were normalized to relative abundances (%), with the total sum of each sample scaled to 100% to ensure comparability. Visualization of the microbial compositions was generated using the ggplot2 package in R.

### Metabolite Analysis

Plants were subjected to 16SC VOC or DBC treatments for 10 days. Three whole seedlings were transferred using sterile forceps into 12-well plates containing 1 mL of sterile Milli-Q water per well and incubated for 2 hours. The of water was collected in Eppendorf tubes and flash-frozen in liquid nitrogen. Additionally, the roots and shoots were collected for further metabolite analysis.

Plant metabolites were determined as in Türksoy et al., (2025) (Türksoy et al., 2025). Roots and shoots (five biological replicates per condition) were extracted by incubation with 0.5 ml of a pre-cooled methyl-*tert*-butyl ether/methanol mixture (MTBE:MeOH, 3:1, v:v), spiked with 10 μg ml^−1^ [^13^C] L-valine, following Berková et al., (2024) (Berková et al., 2024). 100 µl extract aliquotes were dried, derivatized by 10 μl of methoximation solution (40 mg of methoxyamine hydrochloride in 1 ml of pyridine), and incubated for 90 min at 30 °C with continuous shaking. After the incubation, 40 μl of silylation solution (*N*-methyl-*N*-trimethylsilyltrifluoroacetamide) was added, and the mixture was incubated for 30 min at 37 °C with continuous shaking.

Exudates were collected as desribed by Türksoy et al., (2025) (Türksoy et al., 2025), spiked with [^13^C] L-valine for internal calibration and processed in the same way.

For the GC-MS (Gas Chromatography Mass Spectrometry) analysis, the derivatized samples were injected onto the Zebron ZB-5MS plus GC column (Phenomenex; 30 m × 0.25 mm × 0.25 μm; 10 m guard column). Helium was used as the carrier gas at a flow rate of 1.2 ml min⁻¹. Metabolites were separated using the following temperature program: 70 °C for 5 min, ramped at 10 °C min⁻¹ to 320 °C, and held at 320 °C for 5 min. The analytes were ionized using electron ionization mode with an electron energy of 70 eV and an emission current of 50 μA. The transfer line and ion source temperatures were set to 250 °C. Data were acquired in full-scan mode over an m/z range of 50–750.

Data were processed using Compound Discoverer 3.3 (Thermo; peak detection settings: mass tolerance 5 ppm, TIC threshold 25,000, and S/N threshold 3) and searched against the NIST 2023 library, the GC-Orbitrap Metabolomics library, and an in-house library. Only metabolites that met stringent identification criteria, including a spectral match score > 80 and ΔRI < 1%, were included in the final list of identified compounds. Quantitative differences were validated by manual peak assignment in Skyline 19.1 (Pino et al., 2020) using extracted ion chromatograms with a 5-ppm tolerance.

### Statistical analysis

Data visualization of graphs, heatmaps and statistical analyses were performed using GraphPad Prism (version 11). For multiple comparisons, a two-way ANOVA was utilized, followed by Tukey’s HSD (honestly significant difference) *post hoc* pairwise tests. Data normality was assessed using the Shapiro–Wilk test. In all figures, different letters indicate statistically significant differences between group means (*p* < 0.05). In the box plots, the central bars indicate the median, the lower and upper box limits represent the 25th and 75th percentiles, respectively, and the whiskers indicate the minimum and maximum values. Each dot represents an individual sample. Experiments were conducted with four independent replicates, each consisting of 10 plants (excluding any dead plants). For anion analysis, each data point represents a pooled sample of three plant shoots, analyzed in four independent replicates. All experiments were repeated at least twice to ensure reproducibility. Principal Component Analysis (PCA) was performed in R using the prcomp function to evaluate multivariate differences among treatments. Data were mean-centered and scaled to unit variance prior to analysis, and the first two principal components were visualized using ggplot2.

## Data availability

All original data and materials will be provided on request to the corresponding authors.

## Acknowledgement

We thank Qi Wang from Max Planck Institute for Plant Breeding Research providing *pho2* seeds. The authors declare no competing interests.

## Author Contribution

G.M.T., and S.K. conceptualized the project and designed the experiments. S.K. supervised the project. G.M.T. and J.S. conducted the experiments. G.M.T. analyzed the data. M.B. and M.Č. performed GC–MS and analyzed metabolite profiles. G.M.T. wrote original draft of the manuscript. G.M.T. generated figures. S.K. edited the manuscript. All authors read and approved the final manuscript.

## Supplementary Data

**Figure S1.**
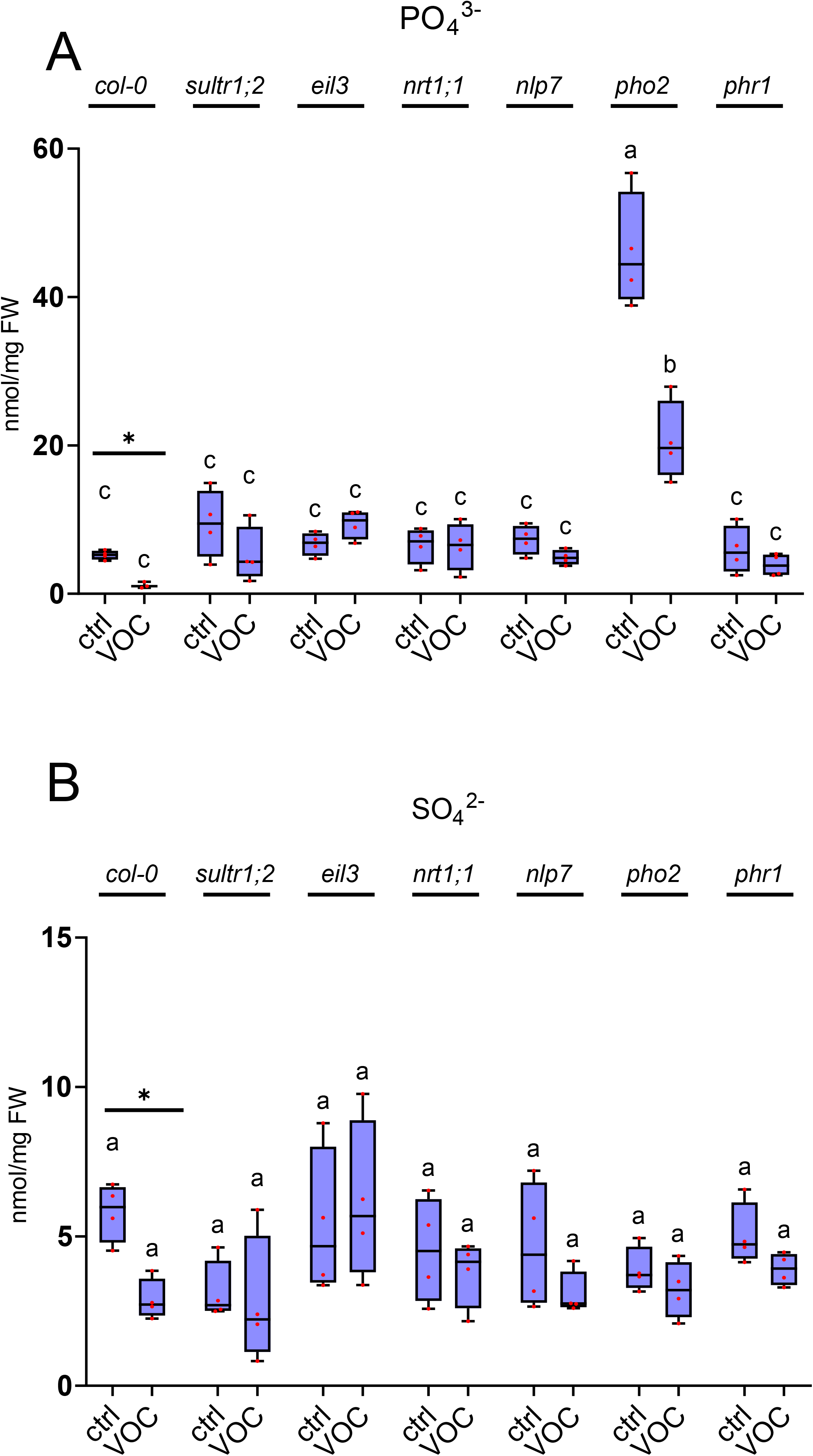
Shoot anion levels upon 10-day 16SC VOC exposure. Shoot **A)** phosphate and **B)** sulfate levels of WT, sultr1;2, eil3, nrt1;1, nlp7, pho2, and phr1 plants following 10 days of 16SC DBC treatment. Letters indicate statistically significant differences between each means(two-way ANOVA and Tukey’s post-hoc test, p < 0.05, A) genotype: F(6, 41) = 71.60, p <0.0001, treatment: F(1, 41) = 29.49, p < 0.0001; interaction: F(6, 41) = 14.38, p < 0.0001. B)genotype: F(6, 42) = 2.576, p = 0.0324), treatment: F(1, 42) = 5.377, p = 0.0253, interaction: F(6, 42) = 1.090, p = 0.3844 n = 4; each dot represents pooled anions from three plants).Asterisks (*) indicate pairwise comparison of two groups (Student’s t-test, p < 0.05).

**Figure S2.**
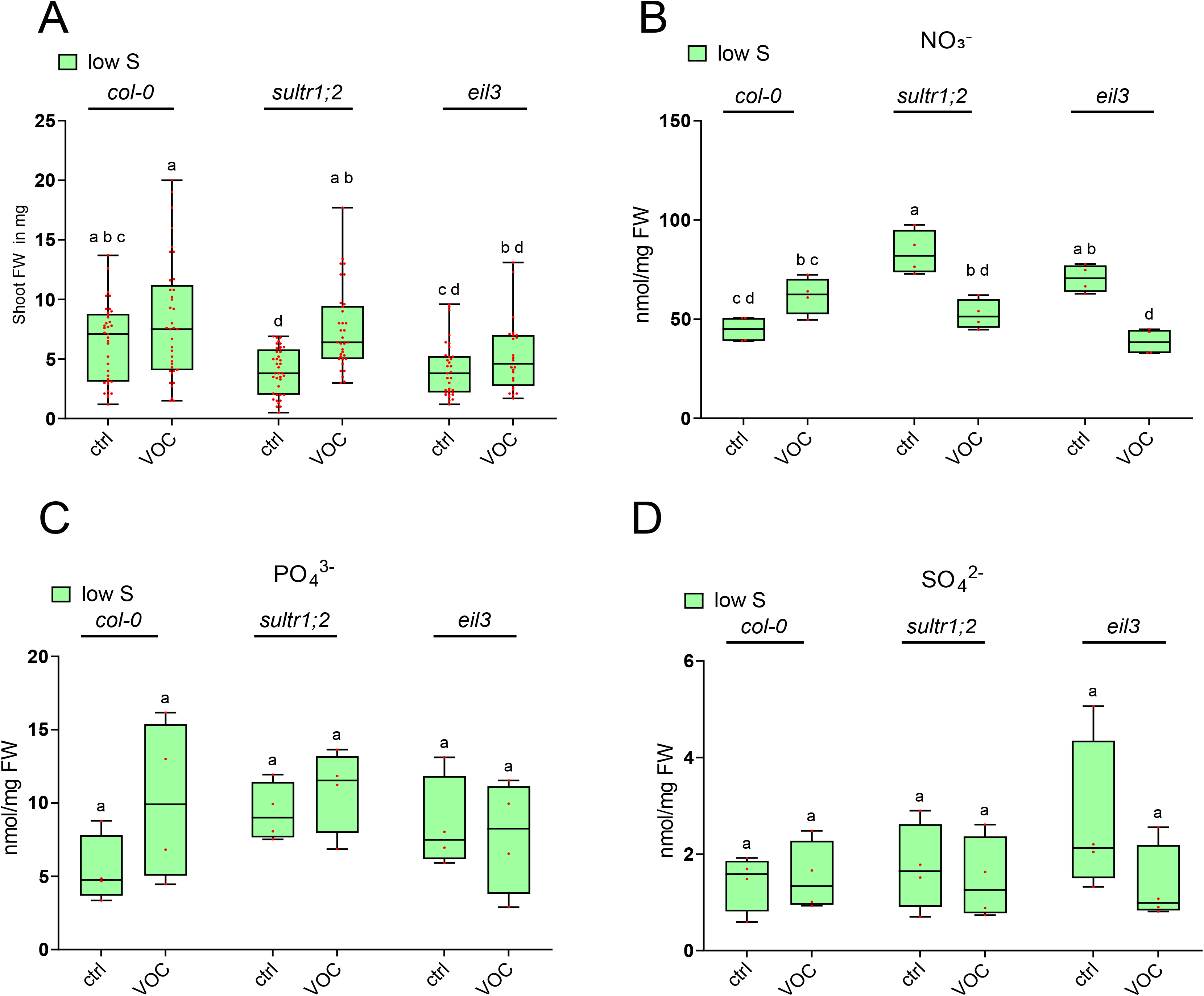
Possible involvement of nutrient pathway genes in 16SC VOC-mediated growth modulation of Arabidopsis thaliana under S limitation. **A)** Fresh weight (mg) of WT, sultr1;2, and eil3 seedlings under sulfur deficiency after 10 days of 16SC VOC treatment. Letters indicate significant differences between each means (two-way ANOVA and Tukey’s post-hoc test, p < 0.05, genotype: F(1, 198) = 20.64, p < 0.0001), treatment: F (2, 198) = 9.843, p < 0.0001, interaction: F(2, 198) = 2.132, p = 0.1213). n = 40; 4 independent replicates with 10 seedlings each). Shoot **B)** nitrate, **C)** phosphate, and **D)** sulfate levels of WT, sultr1;2, and eil3 plants under sulfur deficiency. Letters indicate significant differences between each means (two-way ANOVA and Tukey’s post-hoc test, p < 0.05, B) genotype: F(1, 18) = 21.06, p = 0.0002, treatment: F(2, 18) = 7.809, p = 0.0036; inter action: F(2, 18) = 23.17, p < 0.0001 C) genotype: F(2, 18) = 1.089, p = 0.3577, treatment: F(1, 18) = 1.654, p = 0.2147, interaction: F(2, 18) = 1.255,p = 0.3088. D) genotype: F(2, 18) = 0.6248, p = 0.5466, treatment: F(1, 18) =1.511, p = 0.2349, interaction: F(2, 18) = 1.131, p = 0.3447, n = 4; each dot represents pooled anions from three plants).

**Figure S3.**
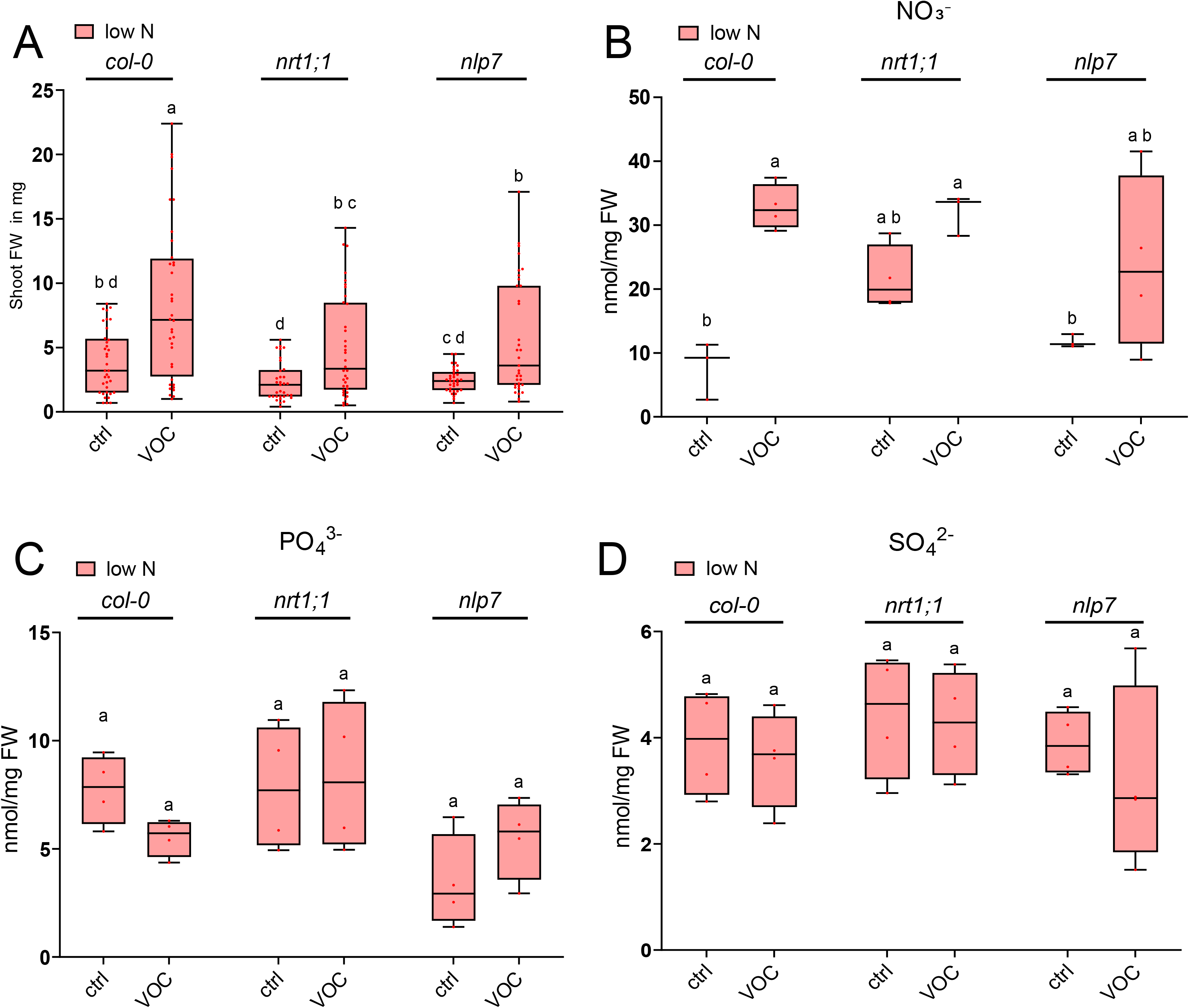
Possible involvement of nutrient pathway genes in 16SC VOC-mediated growth modulation of Arabidopsis thaliana under N limitation. **A)** Fresh weight (mg) of WT, nrt1;1, and nlp7 seedlings under nitrate deficiency after 10 days of 16SC VOC treatment. Letters indicate significant differences between each means (two-way ANOVA and Tukey’s post-hoc test, p < 0.05, genotype: F(1, 205) = 45.69,p<0.0001,treatment: F(2, 205) = 9.178, p = 0.0002, interaction: F(2, 205) = 1.259, p = 0.2862, n = 40; 4 independent replicates with 10 seedlings each). Shoot **B)** nitrate, **C)** phosphate, and **D)** sulfate levels of WT, nrt1;1, and nlp7 plants under nitrate deficiency. Letters indicate significant differences between each group means (two-way ANOVA and Tukey’s post-hoc test, p < 0.05, B) genotype: F(1, 15) = 26.27, p = 0.0001, treatment: F(2, 15) = 2.949, p = 0.0832, interaction: F(2, 15) = 2.214, p = 0.1437 C) genotype: F(2, 18) = 5.054, p = 0.0181, treatment: F(1, 18) = 0.01630, p = 0.8998, interaction: F(2, 18) = 1.761, p = 0.2002 D) genotype: F(2, 18) = 1.055, p = 0.3687, treatment: F(1, 18) = 0.657, p = 0.4280; interaction:F(2, 18) = 0.1083, p = 0.8980, n = 4; each dot represents pooled anions from three plants).

**Figure S4.**
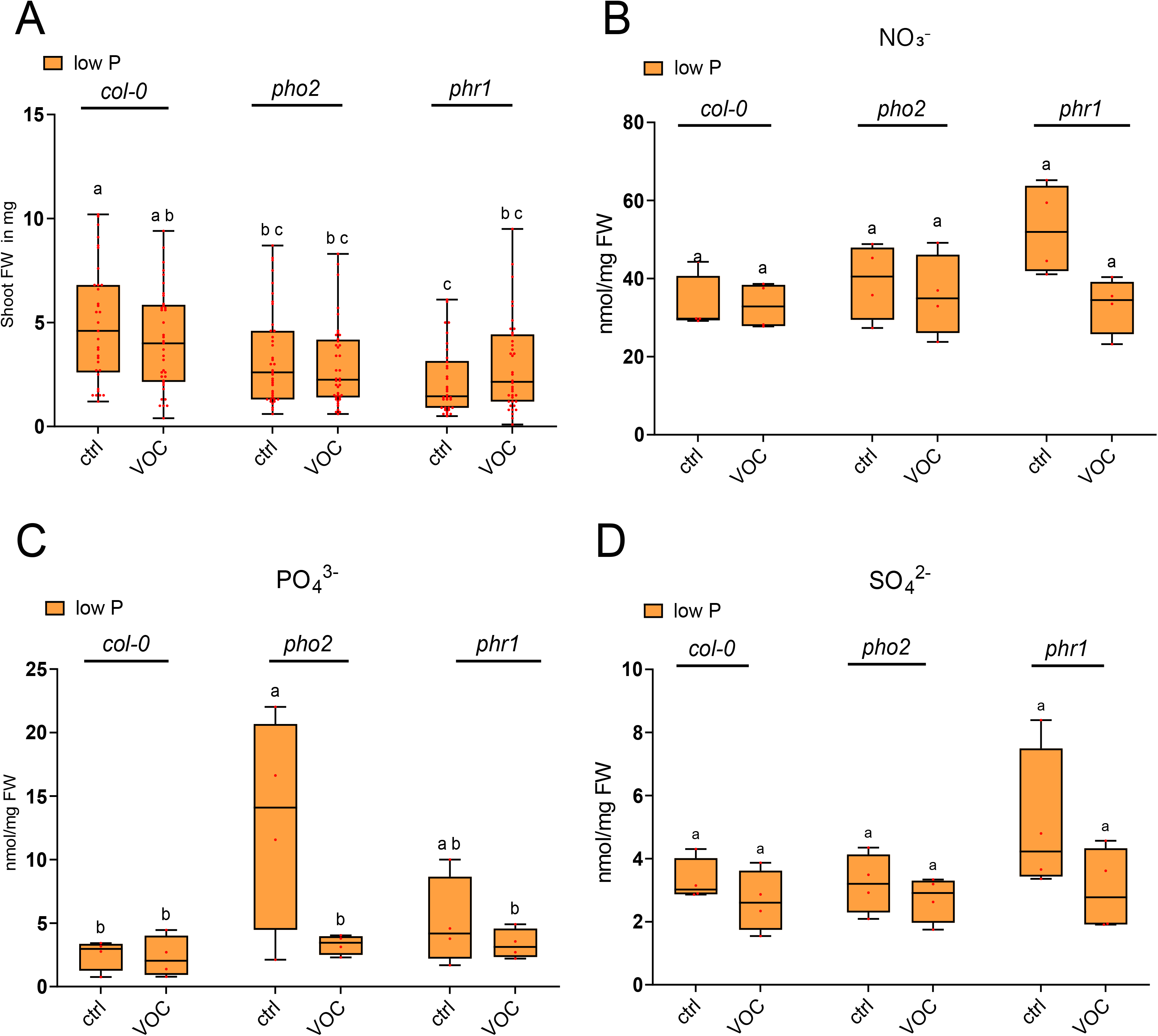
Possible involvement of nutrient pathway genes in 16SC VOC-mediated growth modulation of Arabidopsis thaliana under P limitation. **A)** Fresh weight (mg) of 10 days 16 SC VOCs treated Arabidopsis thaliana seedlings WT, pho2 and phr1 under phosphorus deficiency for 10 days. Letters indicates statistically significant differences between each means, (two-way ANOVA and Tukey’s post-hoc test, p < 0.05,genotype: F(2, 215) = 14.60, p < 0.0001, treatment: F(1, 215) = 0.2536, p = 0.6151, interaction: F(2, 215) = 2.292, p = 0.1035, n = 40; 4 independent replicates with 10 seedlings each) Shoot **B)** nitrate **C)** phosphate **D)** sulfate levels of 10 days 16SC VOC treated WT, pho2 and phr1 plants under phosphorus deficiency Letters indicates statistically significant differences between each means, (two-way ANOVA and Tukey’s post-hoc test (p<0.05), B) genotype: (1, 18) = 4.489, p = 0.0483, treatment: F(2, 18) = 2.371, p = 0.1219, interaction: F(2, 18) = 2.625, p = 0.0999 C) genotype: F(2, 18) = 4.647, p = 0.0236, treatment: F(1, 18) = 5.996, p = 0.0248, interaction: F(2, 18) = 3.528, p = 0.0510. D) genotype: F(1, 18) = 4.078, p = 0.0586, treatment: F(2, 18) = 1.799, p = 0.1940; interaction: F(2, 18) = 0.8951, p = 0.4260, n=4 each dot represent pooled three plants anion levels).

**Figure S5.**
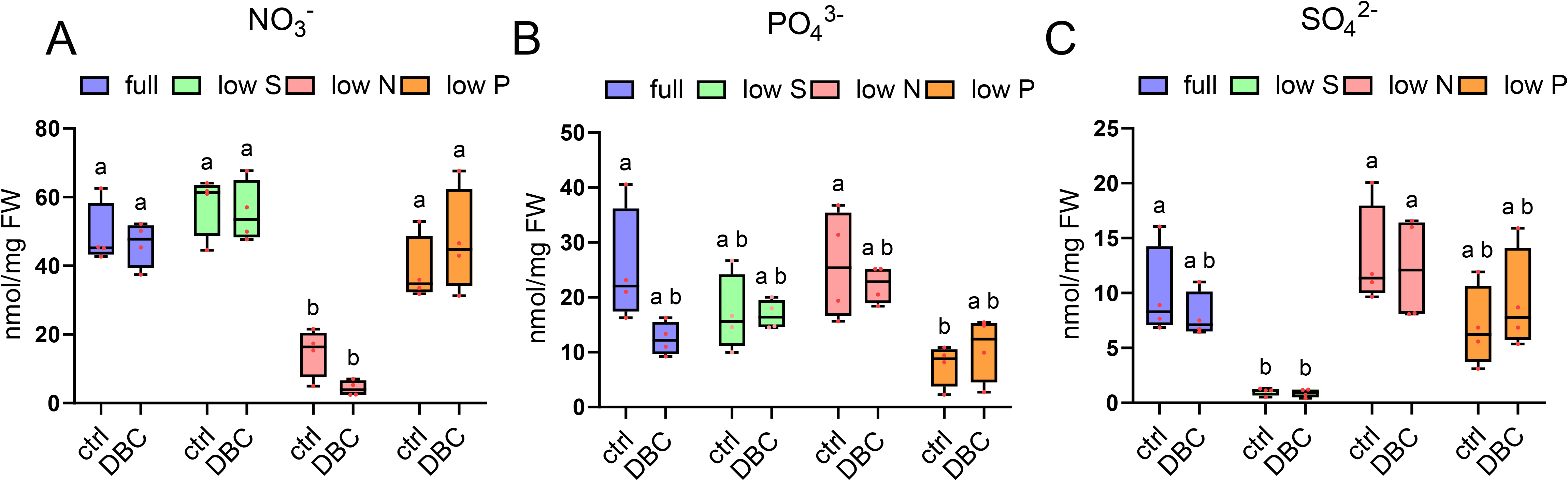
Effect of nutrient limitation on shoot anion levels upon 10-day 16SC DBC exposure. Shoot **A)** nitrate **B)** phosphate **C)** sulfate level of 10 days 16SC DBC treated Arabidopsis thaliana seedlings under sulfur, nitrate and phosphorus deficiencies for 10 days. Letters indicates statistically significant differences between each means, two-way ANOVA and Tukey’s post-hoc test (p<0.05), A)media: F(3, 24) = 40.62, p < 0.0001, treatment: F(1, 24) = 0.2829, p = 0.5997, interaction: F(3, 24) = 1.464, p = 0.2493 B) media: F(3, 24) = 7.178, p = 0.0013, treatment: F(1, 24) = 2.113, p = 0.1590, interaction:F(3, 24) = 2.215, p = 0.1124 C) media: F(3, 24) = 14.93, p < 0.0001, treatment: F(1, 24) = 0.01710, p = 0.8970, interaction: F(3, 24) = 0.5216, p = 0.6715, n=40 4 technical replicates in each 10 seedlings.

**Figure S6.**
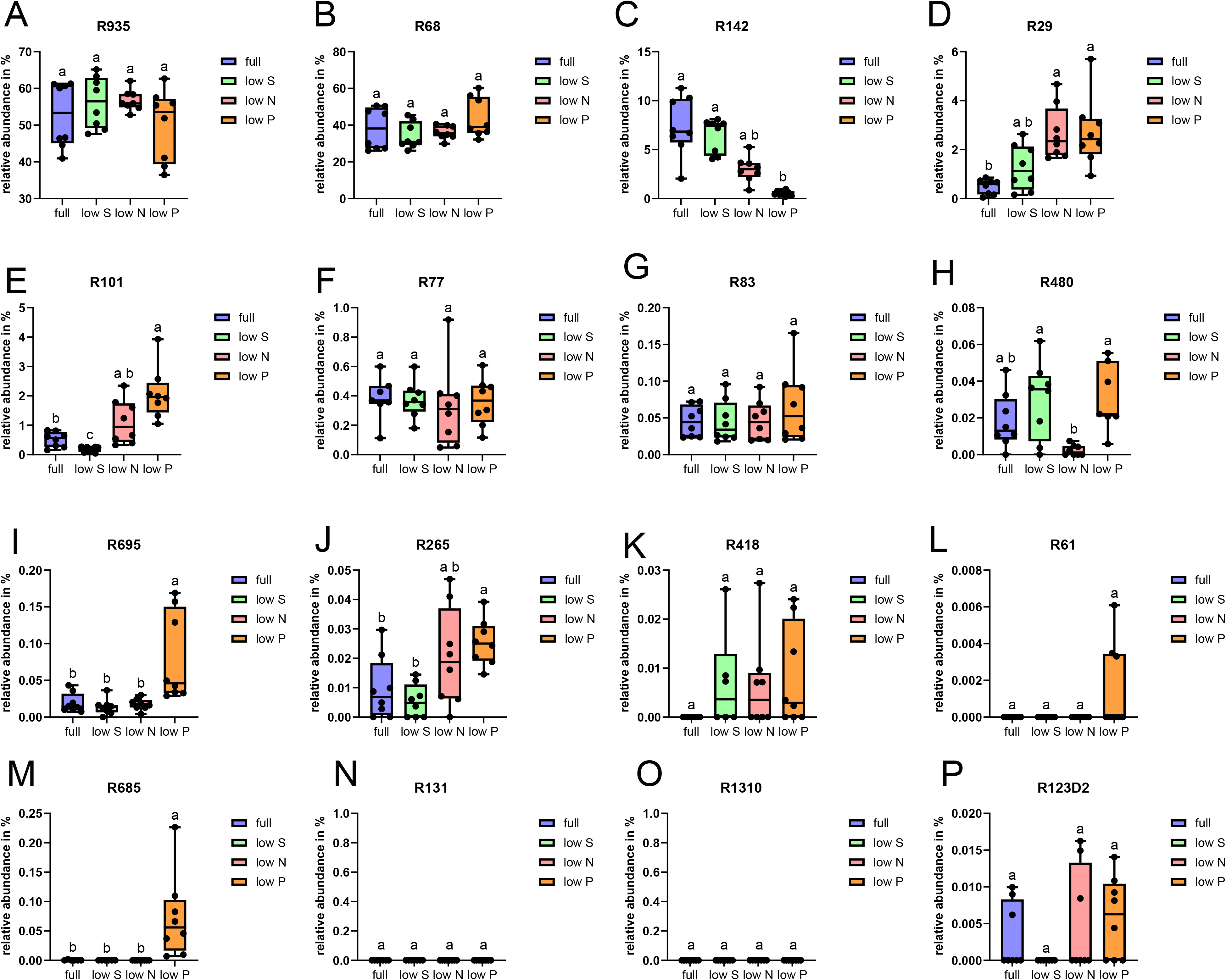
Effect of nutrient limitation on colonized individual strain relative abundance Box plots showing individual strains. **A)** R935 **B)** R68 **C)** R142 **D)** R29 **E)** R101 **F)** R77 **G)** R83 **H)** R480 **I)** R695 **J)** R265 **K)** R418 **L)** R61 **M)** R685 **N)** R131 **O)** R1310 **P)** R123D2 relative abundance (%) colonizing Arabidopsis thaliana roots under full media, low sulfur, low nitrate, and low phosphate conditions for 10 days. Letters indicates statistically significant differences between each group means (Kruskal–Wallis test followed by Dunnett’s post hoc test, p < 0.05), n=8, outliers identified by the ROUT method (Q = 1%) were excluded from the analysis.)

**Figure S7.**
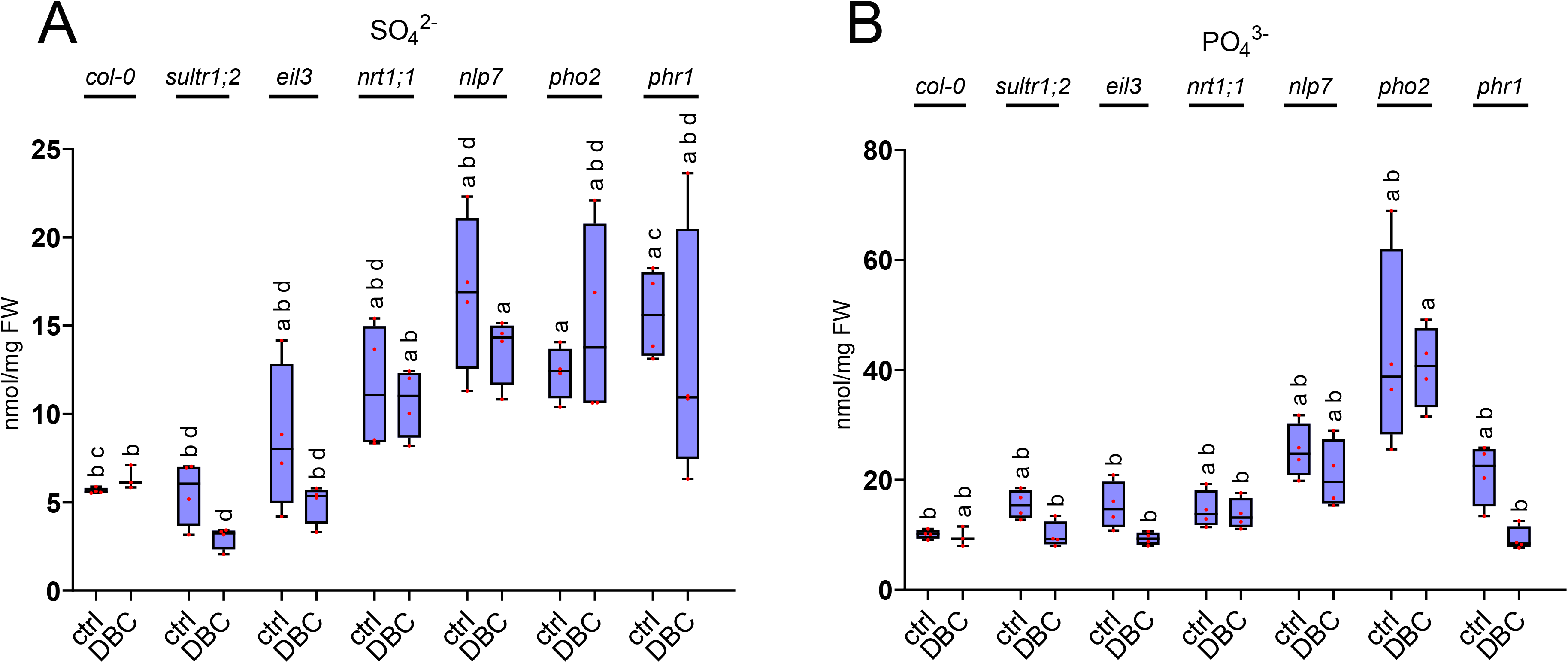
Shoot anion levels upon 10-day 16SC DBC exposure Shoot. **A**) sulfate and **B)** phosphate levels of 10 days 16SC DBC treated WT, sultr1;2, eil3, nrt;1, nlp7, pho2 and phr1 plants were measured. Letters indicate statistically significant differences between each means, (two-way ANOVA and Tukey’s post-hoc test (p>0.05), A) genotype: F(6, 41) = 13.49, p < 0.0001, treatment: F(1, 41) = 2.173, p = 0.1481, interaction: F(6, 41) = 0.9373, p = 0.4790 B) genotype: F(6, 41) = 24.01,p < 0.0001), treatment: F(1, 41) = 6.851, p = 0.0124, interaction: F(6, 41) = 0.7325, p = 0.6262, n=4 each dot represent pooled three plants anion levels

**Figure S8.**
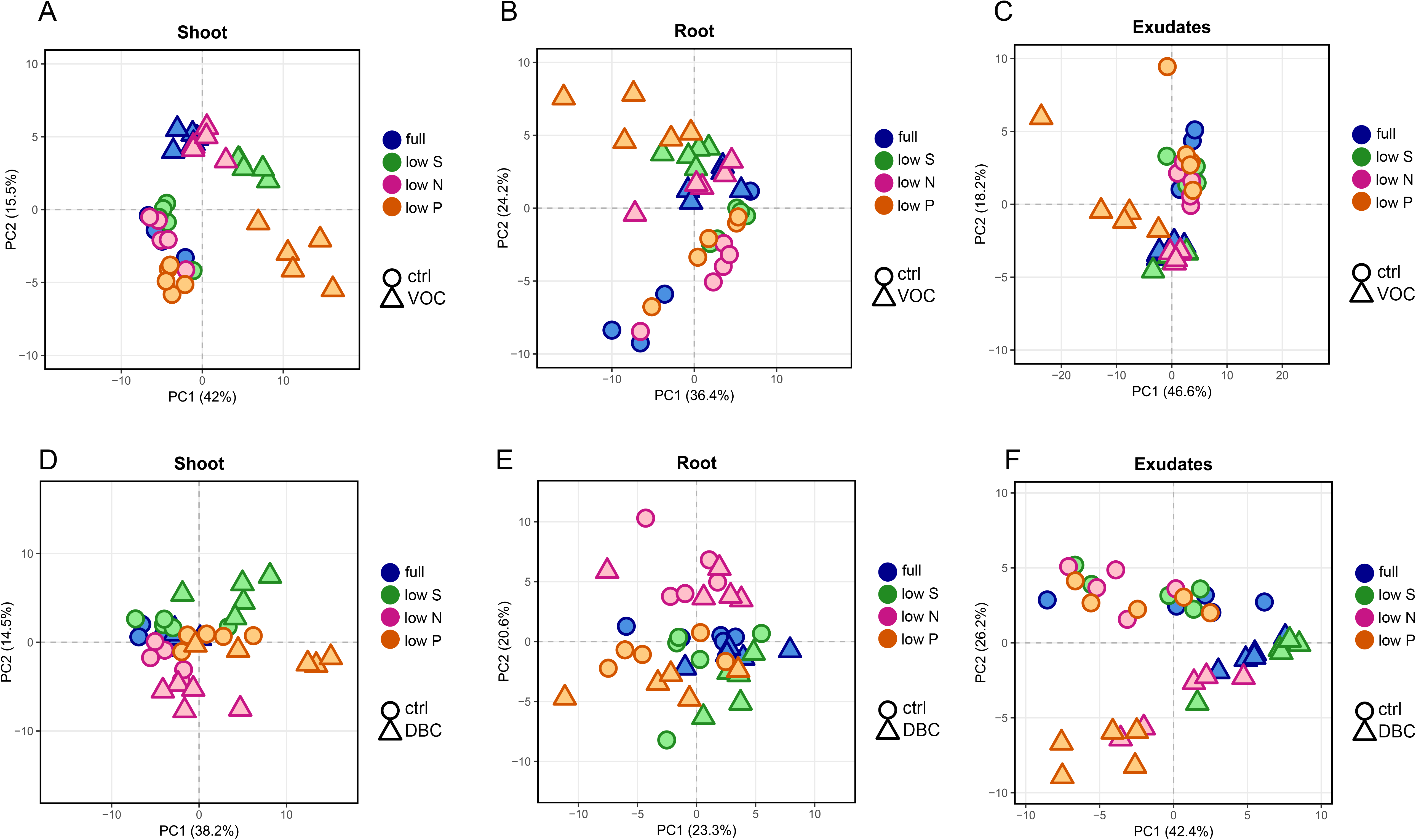
Plant compartments and exudate metabolite difference between VOC and DBC treatments under deficiencies. Principal coordinates analysis (PCoA) showing the effects of nutrient deficiencies under VOC versus mock treatment in the metabolite profiles of **A)** shoots, **B)** roots, and **C)** exudates, and under DBC versus mock treatment in the metabolite profiles of **D)** shoots, **E)** roots, and **F)** exudates.

**Figure S9.**
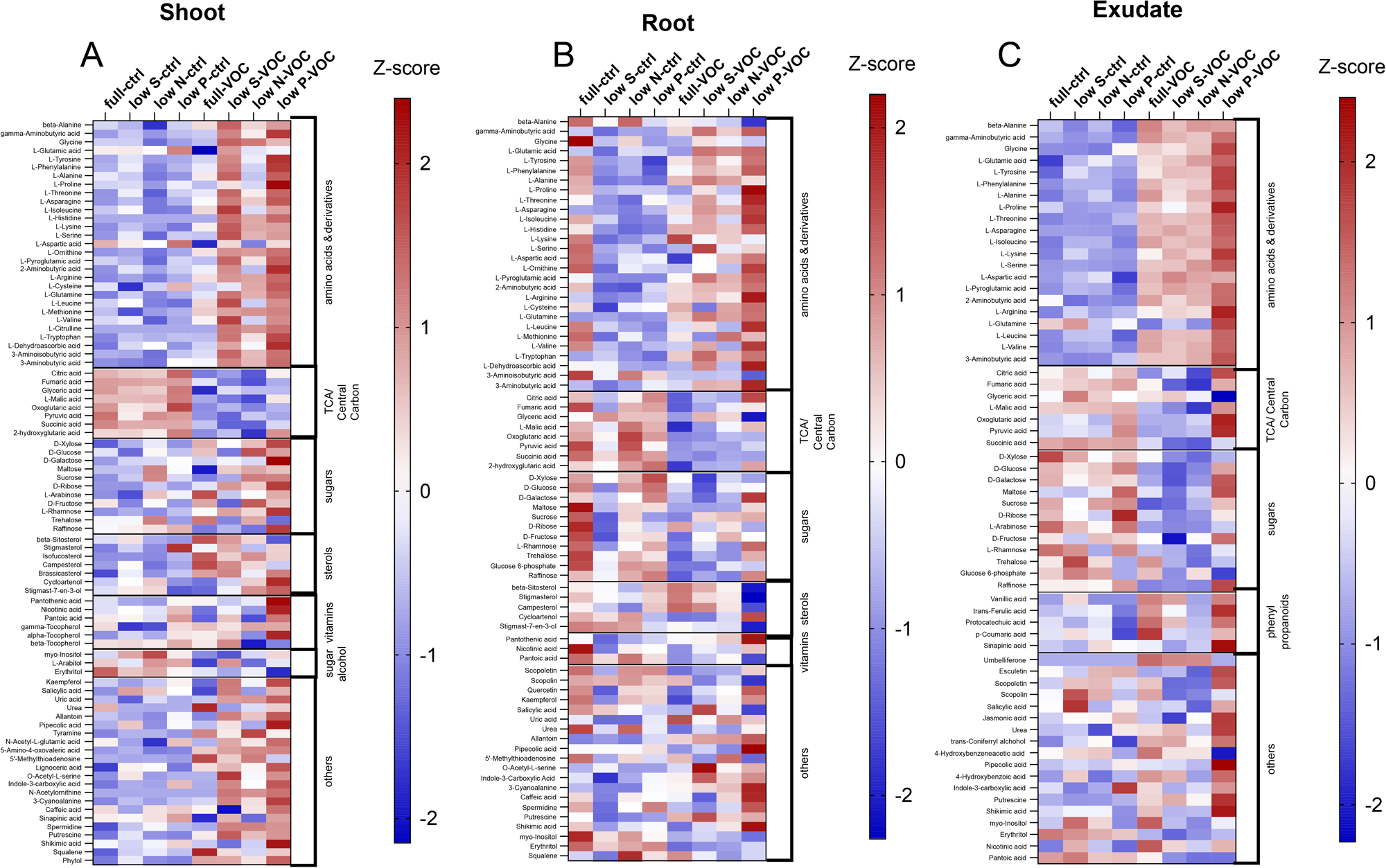
Plant compartments and exudate metabolite difference between VOC and mock treatments under deficiencies. Heatmaps showing metabolite changes in **A)** roots,**B)** shoots, and **C)** exudates under full medium and sulfur, nitrogen, and phosphorus deficienciesupon mock and VOC treatment.

**Figure S10.**
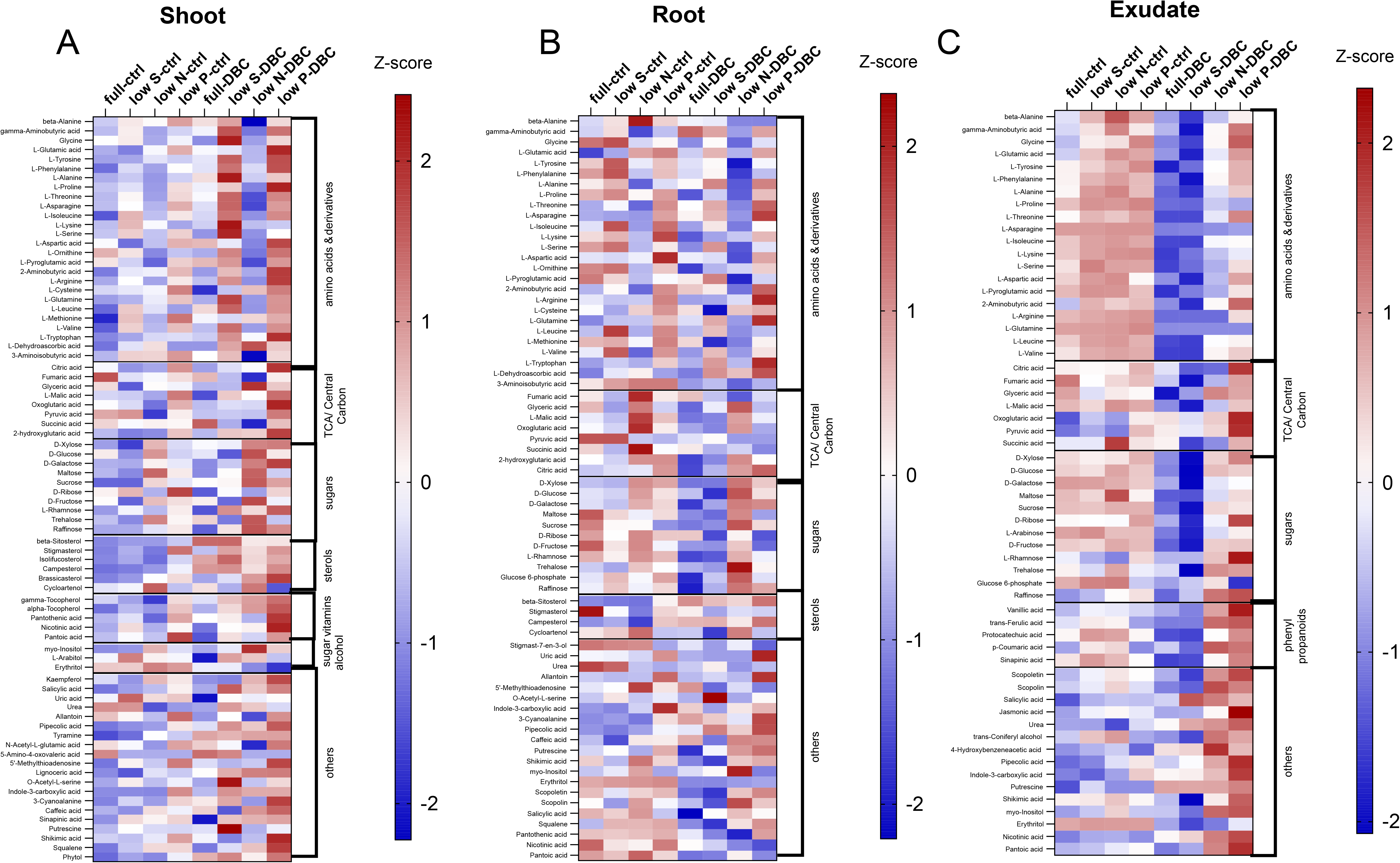
Plant compartments and exudate metabolite difference between DBC and mock treatments under deficiencies. Heatmaps showing metabolite changes in **A)** roots, **B)** shoots, and **C)** exudates under full medium and sulfur, nitrogen, and phosphorus deficiencies upon mock and DBC treatment.upon mock and DBC treatment.

**Table S1.** Composition of nutrient solutions.

| Compound | Concentration full media | Concentration sulfate limitation | Concentration nitrate limitation | Concentration phosphate limitation |
| --- | --- | --- | --- | --- |
| Macroelements |  |  |  |  |
| Ca(NO<sub>3</sub>)<sub>2</sub> x 4H<sub>2</sub>O | 1.5 mM | 1.5 mM |  | 1.5 mM |
| KNO<sub>3</sub> | 1 mM | 1 mM | 0.8 mM | 1 mM |
| KH<sub>2</sub>PO<sub>4</sub> | 0.75 mM | 0.75 mM | 0.75 mM | 0.072 mM |
| MgSO<sub>4</sub> x 7H<sub>2</sub>O | 0.75 mM | 0.0225 mM | 0.75 mM | 0.75 mM |
| Fe-EDTA | 0.1 mM | 0.1 mM | 0.1 mM | 0.1 mM |
| MgCl<sub>2</sub> x 6H<sub>2</sub>O |  | 0.75 mM |  |  |
| CaCl<sub>2</sub> |  |  | 1.5 mM |  |
| KCl |  |  |  | 0.69 mM |
| Microelements | Concentrations |  |  |  |
| MnCl<sub>2</sub>x4H<sub>2</sub>O | 10 µM |  |  |  |
| H<sub>3</sub>BO<sub>3</sub> | 50 µM |  |  |  |
| ZnCl<sub>2</sub> | 1.75 µM |  |  |  |
| CuCl<sub>2</sub> | 0.5 µM |  |  |  |
| Na<sub>2</sub>MoO<sub>4</sub> | 0.8 µM |  |  |  |
| KI | 1 µM |  |  |  |
| CoCl<sub>2</sub>x6H<sub>2</sub>O | 0.1 µM |  |  |  |

**Table S2.**
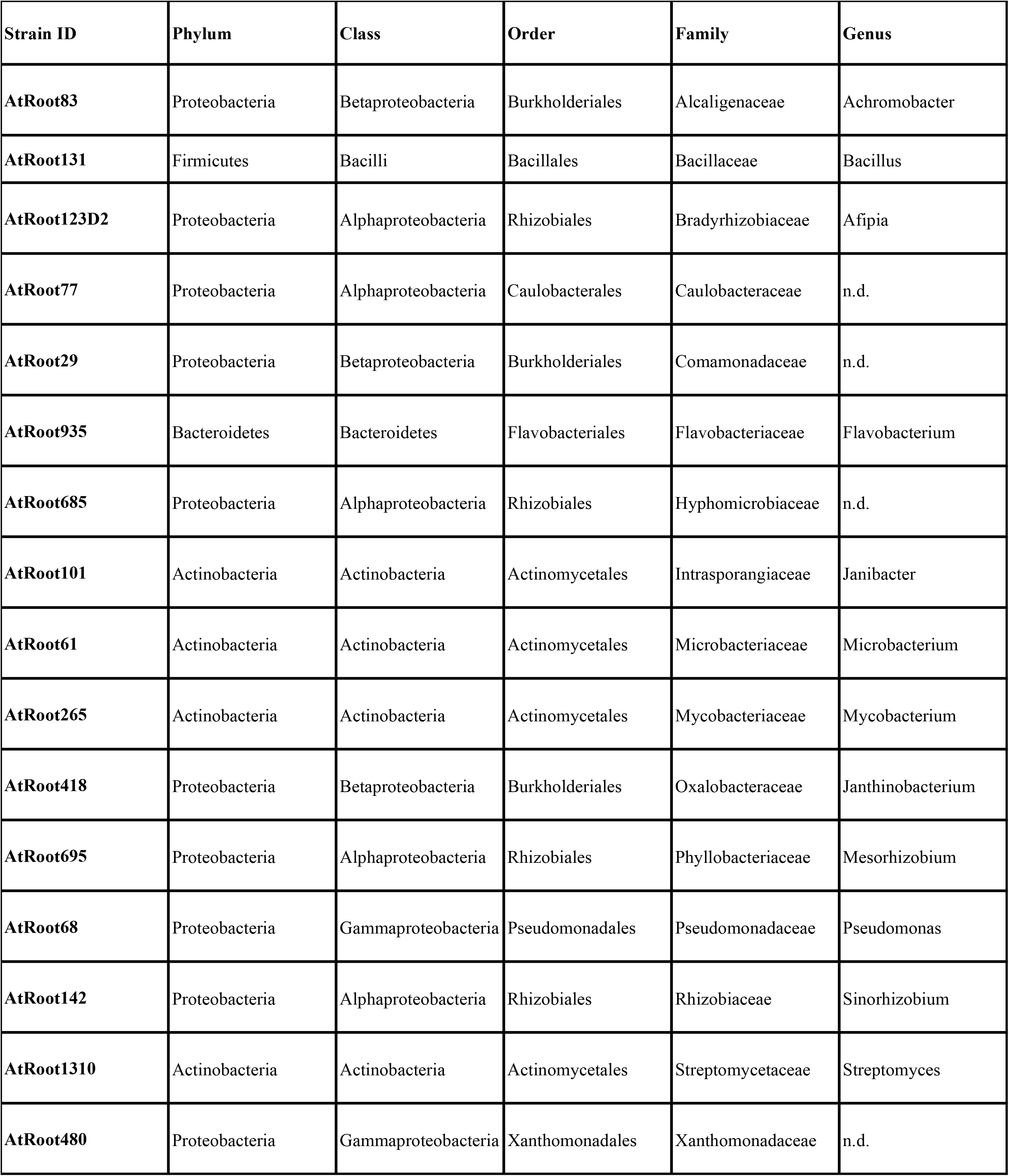
List of 16SC members.

**Table S3.**
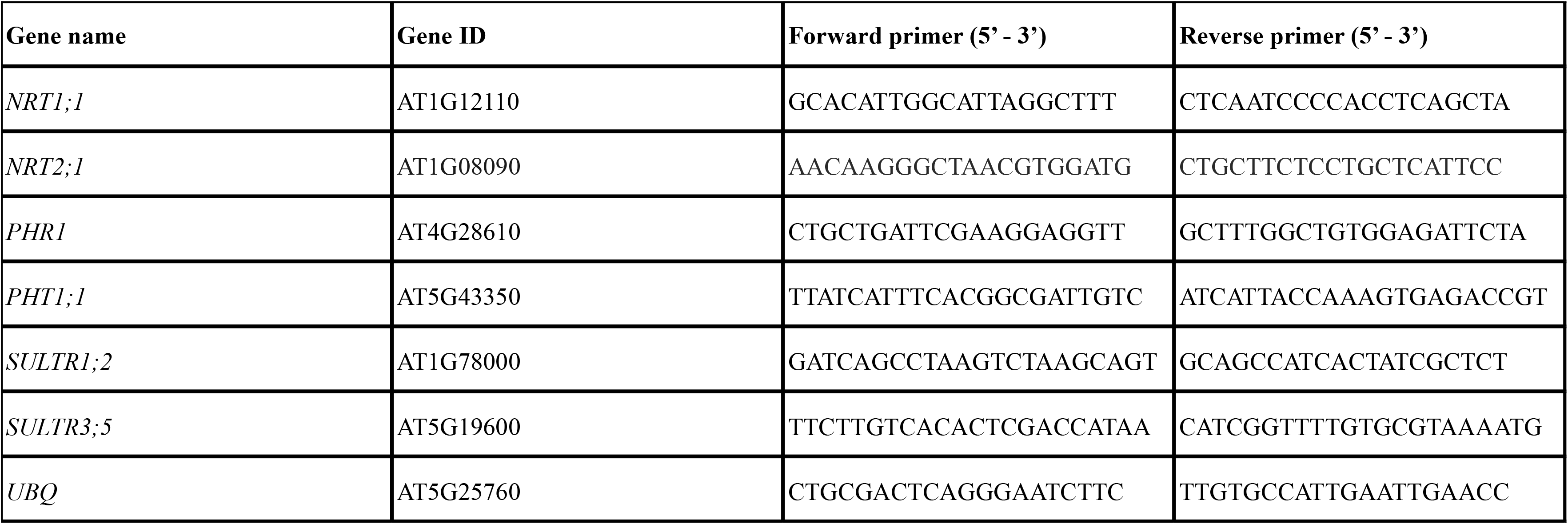
Primers used in qPCR.

## References

Allard-Massicotte, R., Tessier, L., Lécuyer, F., Lakshmanan, V., Lucier, J. F., Garneau, D., Caudwell, L., Vlamakis, H., Bais, H. P., & Beauregard, P. B. (2016). Bacillus subtilis early colonization of Arabidopsis thaliana roots involves multiple chemotaxis receptors. MBio, 7(6). https://doi.org/10.1128/MBIO.01664-16;PAGEGROUP:STRING:PUBLICATION

Almeida, O. A. C., de Araujo, N. O., Mulato, A. T. N., Persinoti, G. F., Sforça, M. L., Calderan-Rodrigues, M. J., & Oliveira, J. V. de C. (2023). Bacterial volatile organic compounds (VOCs) promote growth and induce metabolic changes in rice. Frontiers in Plant Science, 13. 10.3389/fpls.2022.1056082

Aung, K., Lin, S. I., Wu, C. C., Huang, Y. T., Su, C. L., & Chiou, T. J. (2006). pho2, a phosphate overaccumulator, is caused by a nonsense mutation in a microRNA399 target gene. Plant Physiology, 141(3), 1000–1011. 10.1104/PP.106.078063

Aziz, M., Nadipalli, R. K., Xie, X., Sun, Y., Surowiec, K., Zhang, J. L., & Paré, P. W. (2016). Augmenting sulfur metabolism and herbivore defense in arabidopsis by bacterial volatile signaling. Frontiers in Plant Science, 7(APR2016), 182012. 10.3389/FPLS.2016.00458/BIBTEX

Bailly, A., Groenhagen, U., Schulz, S., Geisler, M., Eberl, L., & Weisskopf, L. (2014). The inter-kingdom volatile signal indole promotes root development by interfering with auxin signalling. Plant Journal, 80(5), 758–771. 10.1111/tpj.12666

Berková, V., Berka, M., Štěpánková, L., Kováč, J., Auer, S., Menšíková, S., Ďurkovič, J., Kopřiva, S., Ludwig-Müller, J., Brzobohatý, B., & Černý, M. (2024). The fungus Acremonium alternatum enhances salt stress tolerance by regulating host redox homeostasis and phytohormone signaling. Physiologia Plantarum, 176(3). 10.1111/ppl.14328

Bulgarelli, D., Schlaeppi, K., Spaepen, S., Ver, E., Van Themaat, L., & Schulze-Lefert, P. (2013). Structure and Functions of the Bacterial Microbiota of Plants. 10.1146/annurev-arplant-050312-120106

Bustos, R., Castrillo, G., Linhares, F., Puga, M. I., Rubio, V., Pérez-Pérez, J., Solano, R., Leyva, A., & Paz-Ares, J. (2010). A Central Regulatory System Largely Controls Transcriptional Activation and Repression Responses to Phosphate Starvation in Arabidopsis. PLOS Genetics, 6(9), e1001102. 10.1371/JOURNAL.PGEN.1001102

Callahan, B. J., McMurdie, P. J., Rosen, M. J., Han, A. W., Johnson, A. J. A., & Holmes, S. P. (2016). DADA2: High-resolution sample inference from Illumina amplicon data. Nature Methods, 13(7), 581–583. 10.1038/NMETH.3869

Castrillo, G., Teixeira, P. J. P. L., Paredes, S. H., Law, T. F., De Lorenzo, L., Feltcher, M. E., Finkel, O. M., Breakfield, N. W., Mieczkowski, P., Jones, C. D., Paz-Ares, J., & Dangl, J. L. (2017). Root microbiota drive direct integration of phosphate stress and immunity. Nature, 543(7646), 513–518. 10.1038/nature21417

Chai, Y. N., Qi, Y., Goren, E., Chiniquy, D., Sheflin, A. M., Tringe, S. G., Prenni, J. E., Liu, P., & Schachtman, D. P. (2024). Root-associated bacterial communities and root metabolite composition are linked to nitrogen use efficiency in sorghum. MSystems, 9(1). 10.1128/msystems.01190-23

Chen, Y., Li, Y., Fu, Y., Jia, L., Li, L., Xu, Z., Zhang, N., Liu, Y., Fan, X., Xuan, W., Xu, G., & Zhang, R. (2024). The beneficial rhizobacterium Bacillus velezensis SQR9 regulates plant nitrogen uptake via an endogenous signaling pathway. Journal of Experimental Botany, 75(11), 3388–3400. 10.1093/JXB/ERAE125

Cheng, X., Etalo, D. W., van de Mortel, J. E., Dekkers, E., Nguyen, L., Medema, M. H., & Raaijmakers, J. M. (2017). Genome-wide analysis of bacterial determinants of plant growth promotion and induced systemic resistance by Pseudomonas fluorescens. Environmental Microbiology, 19(11), 4638–4656. 10.1111/1462-2920.13927

Cole, B. J., Feltcher, M. E., Waters, R. J., Wetmore, K. M., Mucyn, T. S., Ryan, E. M., Wang, G., Ul-Hasan, S., McDonald, M., Yoshikuni, Y., Malmstrom, R. R., Deutschbauer, A. M., Dangl, J. L., & Visel, A. (2017). Genome-wide identification of bacterial plant colonization genes. PLoS Biology, 15(9). 10.1371/journal.pbio.2002860

del Carmen Orozco-Mosqueda, M., Macías-Rodríguez, L. I., Santoyo, G., Farías-Rodríguez, R., & Valencia-Cantero, E. (2013). Medicago truncatula increases its iron-uptake mechanisms in response to volatile organic compounds produced by Sinorhizobium meliloti. Folia Microbiologica, 58(6), 579–585. 10.1007/s12223-013-0243-9

Elkatmis, B., Türksoy, G. M., Rodríguez, E., Rahmoune, B., Koprivova, A., & Kopriva, S. (2026). Sulfur as a Central Integrator of Plant-Microbe Interactions: From Nutrient Cycling to Immune Signalling and Microbiome Assembly. Journal of Experimental Botany. 10.1093/JXB/ERAG186

Finkel, O. M., Salas-González, I., Castrillo, G., Spaepen, S., Law, T. F., Teixeira, P. J. P. L., Jones, C. D., & Dangl, J. L. (2019). The effects of soil phosphorus content on plant microbiota are driven by the plant phosphate starvation response. PLoS Biology, 17(11). 10.1371/journal.pbio.3000534

Garbeva, P., & Weisskopf, L. (2020). Airborne medicine: bacterial volatiles and their influence on plant health. New Phytologist, 226(1), 32–43. https://doi.org/10.1111/NPH.16282;WGROUP:STRING:PUBLICATION

Griffin, C., Oz, M. T., & Demirer, G. S. (2024). Engineering plant–microbe communication for plant nutrient use efficiency. Current Opinion in Biotechnology, 88, 103150. 10.1016/J.COPBIO.2024.103150

Guo, J. H., Liu, X. J., Zhang, Y., Shen, J. L., Han, W. X., Zhang, W. F., Christie, P., Goulding, K. W. T., Vitousek, P. M., & Zhang, F. S. (2010). Significant acidification in major chinese croplands. Science, 327(5968), 1008–1010. https://doi.org/10.1126/SCIENCE.1182570;WEBSITE:WEBSITE:AAAS-SITE;JOURNAL:JOURNAL:SCIENCE;WGROUP:STRING:PUBLICATION

Gutjahr, C., & Parniske, M. (2013). Cell and developmental biology of arbuscular mycorrhiza symbiosis. Annual Review of Cell and Developmental Biology, 29(Volume 29, 2013), 593–617. https://doi.org/10.1146/ANNUREV-CELLBIO-101512-122413/CITE/REFWORKS

Ho, C. H., Lin, S. H., Hu, H. C., & Tsay, Y. F. (2009). CHL1 Functions as a Nitrate Sensor in Plants. Cell, 138(6), 1184–1194. 10.1016/j.cell.2009.07.004

Hu, L., Robert, C. A. M., Cadot, S., Zhang, X., Ye, M., Li, B., Manzo, D., Chervet, N., Steinger, T., Van Der Heijden, M. G. A., Schlaeppi, K., & Erb, M. (2018). Root exudate metabolites drive plant-soil feedbacks on growth and defense by shaping the rhizosphere microbiota. Nature Communications, 9(1). 10.1038/s41467-018-05122-7

Jacoby, R. P., Koprivova, A., & Kopriva, S. (2021). Pinpointing secondary metabolites that shape the composition and function of the plant microbiome. Journal of Experimental Botany, 72(1), 57–69. 10.1093/JXB/ERAA424

Koprivova, A., Berka, M., Berková, V., Ristova, D., Türksoy, G. M., Schwier, M., Westhoff, P., Cerný, M., Kopriva, S., Brno, I., Zeměd, Z., & Zemědělská, Z. (2025). Multilevel Analysis of Response to Plant Growth-Promoting and Pathogenic Bacteria in Arabidopsis Roots. 10.1094/MPMI-09-25-0125-R, 38(6), 1031–1043. 10.1094/MPMI-09-25-0125-R

Li, N., Li, G., Huang, X., Ma, L., Wang, D., Luo, Y., Cao, X., Zhu, Y., Mu, J., An, R., Zhao, J., Wang, Y., Yang, C., Chen, H., Xu, Y., Jiang, L., Luo, M., Li, X., Dong, Y., … Yu, P. (2026). Large-scale multi-omics unveils host–microbiome interactions driving root development and nitrogen acquisition. Nature Plants 2026 12:2, 12(2), 319–336. 10.1038/s41477-025-02210-7

Liu, X., Zhang, Y., Han, W., Tang, A., Shen, J., Cui, Z., Vitousek, P., Erisman, J. W., Goulding, K., Christie, P., Fangmeier, A., & Zhang, F. (2013). Enhanced nitrogen deposition over China. Nature 2013 494:7438, 494(7438), 459–462. 10.1038/nature11917

Lugtenberg, B., & Kamilova, F. (2009). Plant-growth-promoting rhizobacteria. In Annual Review of Microbiology (Vol. 63, pp. 541–556). 10.1146/annurev.micro.62.081307.162918

Meldau, D. G., Meldau, S., Hoang, L. H., Underberg, S., Wünsche, H., & Baldwin, I. T. (2013). Dimethyl Disulfide Produced by the Naturally Associated Bacterium Bacillus sp B55 Promotes Nicotiana attenuata Growth by Enhancing Sulfur Nutrition. The Plant Cell, 25(7), 2731–2747. 10.1105/TPC.113.114744

Moormann, J., Heinemann, B., & Hildebrandt, T. M. (2022). News about amino acid metabolism in plant–microbe interactions. Trends in Biochemical Sciences, 47(10), 839–850. 10.1016/J.TIBS.2022.07.001

Mukherjee, A., Han, L., Mukhopadhyay, S., Kopriva, S., & Swarup, S. (2025). Sulfur traits in the plant microbiome: implications for sustainable agriculture. Trends in Microbiology, 33(6), 635–649. 10.1016/J.TIM.2025.02.002

Mukherjee, A., Mazumder, M., Verma, A., Tikariha, H., Bhattacharya, R., Ooi, Q. E., & Swarup, S. (2025). A bacterial signal coordinates plant-microbe fitness trade-off to enhance sulfur deficiency tolerance in plants. Cell Host and Microbe, 33(10), 1748–1764.e6. 10.1016/j.chom.2025.09.007

Pieterse, C. M. J., Zamioudis, C., Berendsen, R. L., Weller, D. M., Van Wees, S. C. M., & Bakker, P. A. H. M. (2014). Induced systemic resistance by beneficial microbes. Annual Review of Phytopathology, 52(September), 347–375. 10.1146/annurev-phyto-082712-102340

Pino, L. K., Searle, B. C., Bollinger, J. G., Nunn, B., MacLean, B., & MacCoss, M. J. (2020). The Skyline ecosystem: Informatics for quantitative mass spectrometry proteomics. In Mass Spectrometry Reviews (Vol. 39, Number 3, pp. 229–244). John Wiley and Sons Inc. 10.1002/mas.21540

Rahimlou, S., Bahram, M., & Tedersoo, L. (2021). Phylogenomics reveals the evolution of root nodulating alpha- and beta-Proteobacteria (rhizobia). Microbiological Research, 250, 126788. 10.1016/J.MICRES.2021.126788

Ristova, D., & Kopriva, S. (2022). Sulfur signaling and starvation response in Arabidopsis. IScience, 25(5). 10.1016/J.ISCI.2022.104242

Rizaludin, M. S., Stopnisek, N., Raaijmakers, J. M., & Garbeva, P. (2021). The Chemistry of Stress: Understanding the ‘Cry for Help’ of Plant Roots. Metabolites, 11(6), 357. 10.3390/METABO11060357

Ryu, C. M., Faragt, M. A., Hu, C. H., Reddy, M. S., Wei, H. X., Paré, P. W., & Kloepper, J. W. (2003). Bacterial volatiles promote growth in Arabidopsis. Proceedings of the National Academy of Sciences of the United States of America, 100(8), 4927–4932. 10.1073/pnas.0730845100

Schulz, S., & Dickschat, J. S. (2007). Bacterial volatiles: the smell of small organisms. Natural Product Reports, 24(4), 814–842. 10.1039/B507392H

Shi, J., Wang, X., & Wang, E. (2026). Mycorrhizal Symbiosis in Plant Growth and Stress Adaptation: From Genes to Ecosystems. *Annual* Review of Plant Biology Downloaded from Www.Annualreviews.Org. Guest. 10.1146/annurev-arplant-061722

Tahir, H. A. S., Gu, Q., Wu, H., Raza, W., Hanif, A., Wu, L., Colman, M. V., & Gao, X. (2017). Plant growth promotion by volatile organic compounds produced by Bacillus subtilis SYST2. In Frontiers in Microbiology (Vol. 8, Number FEB). 10.3389/fmicb.2017.00171

Trivedi, P., Leach, J. E., Tringe, S. G., Sa, T., & Singh, B. K. (2020). Plant–microbiome interactions: from community assembly to plant health. Nature Reviews Microbiology 2020 18:11, 18(11), 607–621. 10.1038/s41579-020-0412-1

Tsai, H. H., Tang, Y., Jiang, L., Xu, X., Tendon, V. D., Pang, J., Jia, Y., Wippel, K., Vacheron, J., Keel, C., Andersen, T. G., Geldner, N., & Zhou, F. (2025). Localized glutamine leakage drives the spatial structure of root microbial colonization. Science, 390(6768). 10.1126/science.adu4235

Türksoy, G. M., Berka, M., Wippel, K., Koprivova, A., Carron, R. A., Rüger, L., Černý, M., Andersen, T. G., & Kopriva, S. (2025). Bacterial community-emitted volatiles regulate Arabidopsis growth and root architecture in a distinct manner of those from individual strains. Plant Communications, 6(6), 101351. 10.1016/j.xplc.2025.101351

van der Loo, M. P. J. (2014). The stringdist package for approximate string matching. R Journal, 6(1), 111–122. 10.32614/RJ-2014-011

Wang, W., Shi, J., Xie, Q., Jiang, Y., Yu, N., & Wang, E. (2017). Nutrient Exchange and Regulation in Arbuscular Mycorrhizal Symbiosis. In Molecular Plant (Vol. 10, Number 9, pp. 1147–1158). Cell Press. 10.1016/j.molp.2017.07.012

Weisskopf, L., Schulz, S., & Garbeva, P. (2021). Microbial volatile organic compounds in intra-kingdom and inter-kingdom interactions. Nature Reviews Microbiology, 19(6), 391–404. 10.1038/s41579-020-00508-1

Wippel, K., Tao, K., Niu, Y., Zgadzaj, R., Kiel, N., Guan, R., Dahms, E., Zhang, P., Jensen, D. B., Logemann, E., Radutoiu, S., Schulze-Lefert, P., & Garrido-Oter, R. (2021). Host preference and invasiveness of commensal bacteria in the Lotus and Arabidopsis root microbiota. Nature Microbiology, 6(9), 1150–1162. 10.1038/s41564-021-00941-9

Yang, Y., Pollard, A. M., Höfler, C., Poschet, G., Wirtz, M., Hell, R., & Sourjik, V. (2015). Relation between chemotaxis and consumption of amino acids in bacteria. Molecular Microbiology, 96(6), 1272–1282. 10.1111/MMI.13006

Yoshimoto, N., Takahashi, H., Smith, F. W., Yamaya, T., & Saito, K. (2002). Two distinct high-affinity sulfate transporters with different inducibilities mediate uptake of sulfate in Arabidopsis roots. Plant Journal, 29(4), 465–473. https://doi.org/10.1046/J.0960-7412.2001.01231.X;SUBPAGE:STRING:FULL

Zhang, H., Kim, M. S., Krishnamachari, V., Payton, P., Sun, Y., Grimson, M., Farag, M. A., Ryu, C. M., Allen, R., Melo, I. S., & Paré, P. W. (2007). Rhizobacterial volatile emissions regulate auxin homeostasis and cell expansion in Arabidopsis. Planta, 226(4), 839–851. 10.1007/s00425-007-0530-2

Zhang, H., Sun, Y., Xie, X., Kim, M. S., Dowd, S. E., & Paré, P. W. (2009). A soil bacterium regulates plant acquisition of iron via deficiency-inducible mechanisms. Plant Journal, 58(4), 568–577. 10.1111/j.1365-313X.2009.03803.x

Zhang, J., Liu, Y. X., Zhang, N., Hu, B., Jin, T., Xu, H., Qin, Y., Yan, P., Zhang, X., Guo, X., Hui, J., Cao, S., Wang, X., Wang, C., Wang, H., Qu, B., Fan, G., Yuan, L., Garrido-Oter, R., … Bai, Y. (2019). NRT1.1B is associated with root microbiota composition and nitrogen use in field-grown rice. Nature Biotechnology 2019 37:6, 37(6), 676–684. 10.1038/s41587-019-0104-4

